# Fracking and Ostwald ripening position the lumen of the mouse blastocyst

**DOI:** 10.1101/537985

**Authors:** Julien G. Dumortier, Mathieu Le Verge-Serandour, Anna-Francesca Tortorelli, Annette Mielke, Ludmilla de Plater, Hervé Turlier, Jean-Léon Maître

## Abstract

During mouse preimplantation development, the formation of the blastocoel, a fluid-filled lumen, breaks the radial symmetry of the blastocyst. What controls the formation and positioning of this basolateral lumen remains obscure. We find that accumulation of pressurized fluid fractures cell-cell contacts into hundreds of micron-size lumens. Microlumens eventually discharge their volumes into a single dominant lumen, which we model as a process akin to Ostwald ripening, underlying the coarsening of foams. Using chimeric mutant embryos, we tune the fracking of cell-cell contacts and steer the coarsening of microlumens, allowing us to successfully manipulate the final position of the lumen. We conclude that hydraulic fracture of cell-cell contacts followed by directed coarsening of microlumens sets the first axis of symmetry of the mouse embryo.

## Main text

During preimplantation development, the mammalian embryo forms the blastocyst, which consists of a squamous epithelium, the trophectoderm (TE), enveloping the inner cells mass (ICM) and a fluid-filled lumen, the blastocoel (*1,2*). This lumen forms invariably at the interface between ICM and TE cells and segregates the ICM toward one side of the embryo, hence breaking its radial symmetry. This defines the first symmetry axis in the embryo, which guides the position of the main axes of the mammalian body (*3,4*). The embryonic-abembryonic (Em/Ab) axis designates the axis of the blastocyst, with the ICM and polar TE on the Em side and the blastocoel and mural TE on the Ab side. External constraints, such as the one provided by the zona pellucida, are thought to guide the orientation of the Em/Ab axis (*5, 6*). However, little is known of the mechanisms that are intrinsic to the embryo. Lumen formation and positioning has been studied in the context of apical lumens (*7*) such as the ones formed in vitro by kidney (*8*) or liver (*9, 10*) cells, or in vivo within the epiblast upon implantation of the blastocyst (*11*). Apicobasal polarity allows building an osmotic gradient that draws water from the outside medium into the apical compartment sealed bytightjunctions (*7*). Importantly, the apical membrane is depleted of adhesion molecules and can contain proteins that act as a contact repellent (*12, 13*). This makes the apical membrane weakly adhesive and a favorable interface for the collection of fluid between cells. However, blastocysts are akin to “inverted cysts” (*14, 15*) and form their lumen on the basolateral side of cells, where adhesion molecules concentrate, making this interface mechanically less favorable for fluid to accumulate (*1, 2, 16*) (Fig. S1). Therefore, it is unclear how the blastocoel forms within an adhesive interface and acquires its position between TE and ICM cells.

One possibility would be to separate one or few neighboring TE-ICM contacts and expand the blastocoel from this initial gap (*17*). Alternatively, multiple intercellular gaps could appear between cells, as can be observed between TE cells in electron micrographs of blastocysts from mice (*18*) and non-human primates (*19, 20*), which would merge into a single lumen via unknown mechanisms (21). To investigate this, we used mouse embryos expressing a membrane marker (*22*) to perform high-resolution imaging at the onset of lumen formation, when blastomeres have completed their fifth cleavage (Movie 1). We observed hundreds of micron-size lumens (microlumens), forming simultaneously throughout the embryo between cell-cell contacts and at multicellular interfaces (Fig. 1A). Notably, this includes contacts between ICM cells where the blastocoel never forms (Fig. S2). The size of microlumens evolve rather synchronously throughout the embryo, showing an initial phase of growth followed by a shrinking period (Fig. S2-3). While eventually most microlumens shrink, one continues expanding and becomes the blastocoel (Fig. 1B). From image and data analysis, we characterized two types of microlumens, at either bicellular and multicellular interfaces, and determined stereotypic parameters describing their evolution: for all microlumens but the blastocoel, we identified a swelling phase followed by a discharge phase (Fig. 1B-C, Movie 1). After 87 ± 10 min of steady growth, microlumens between two contacting cells shrink within 68 ± 8 min (Mean± SEM of 35 contacts from 7 embryos). Compared to bicellular microlumens, multicellular microlumens grow 10 times larger over a similar duration (swelling for 75 ± 21 min and discharge for 65 ± 21 min, Mean± SEM from 21 multicellular junctions from 7 embryos), as a result of higher swelling and discharge rates (Fig. 1D-F). In addition, we notice that multicellular microlumens peak with a delay of 30 ± 21 min compared to bicellular microlumens (Fig. 1G). From these observations and analysis, we conclude that the formation of the blastocoel does not result from localized separation of contacts between TE and ICM cells specifically. Instead, transient ruptures of cell-cell contacts result in the formation of myriad lumens throughout the embryo that subsequently disappear, leaving only one remaining.

**Figure 1:**
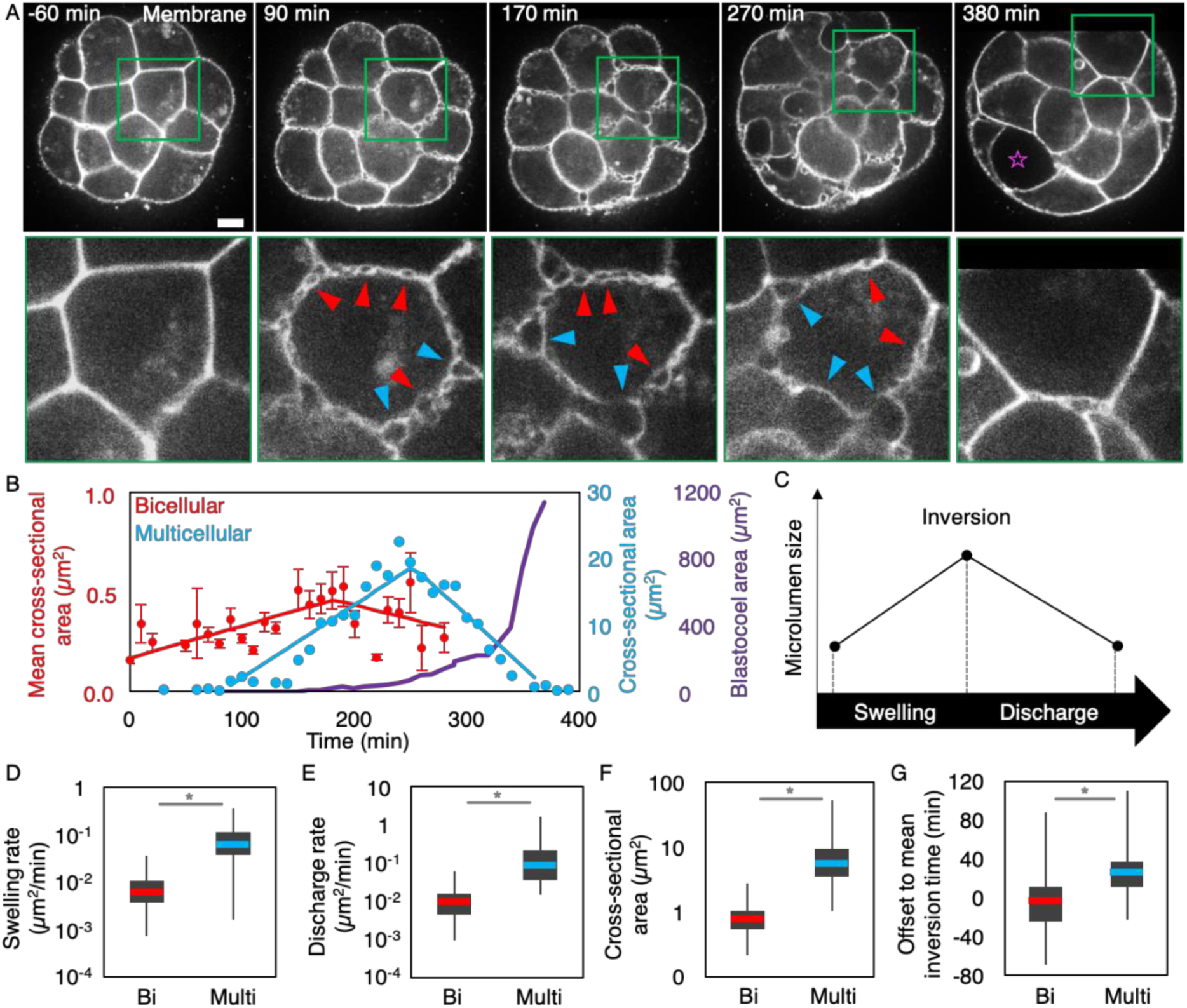
the blastocoel forms at the basolateral interface by swelling and discharge of microlumens. (A) Snapshots of time lapse imaging of blastocoel formation in an embryo with membrane marker (mTmG) from Movie S1. Microlumens form transiently at cell-cell contacts (red arrowheads) and multicellular junctions (blue arrowheads). They swell and then shrink as the blastocoel (magenta star) expands. Green squares are magnified 3 times on the bottom images. Scale bar, 10 *µ*m. (B) Blastocoel (magenta) and microlumens growth dynamics at bicellular (red) or multicellular (blue) junctions. For bicellular microlumens, the mean and SEM of all microlumens at a cell-cell contact are shown. (C) Schematic diagram of the two phases of microlumen lifetime. (D-G) Swelling (D) and shrinkage **(E)** rates, cross-sectional area at the time of inversion (F), and offset to mean inversion time for microlumens at 35 bicellular and 21 multicellular junctions from 7 embryos. (D-F) display data on a log scale. * for Student’s *t* test at p < 10^−2^.

To understand what controls the initiation of the blastocoel, we first investigated how microlumens form at cell-cell contacts. Contacts are enriched in Cdh1, the main cell-cell adhesion molecule of the blastocyst (*23*), which provides them mechanical stability. To visualize the localization of Cdh1 during lumen formation, we generated a transgenic mouse line expressing Cdh1-GFP under its endogenous promoter (*24*) in addition to a membrane marker (*22*). When microlumens form, Cdh1-GFP localization is reorganized, seemingly accumulating at the edges of microlumens (Fig. 2A, Movie 2). This makes the distribution of Cdh1-GFP more heterogeneous (Fig. 28). The reorganization of Cdh1 could directly result from the accumulation of fluid detaching cell-cell contacts. Such hydraulic fracture, or fracking, of cell-cell contacts has been described in vitro when fluid, pressurized at a few hundreds of pascals, is flushed through the basolateral side of mature epithelial monolayers (*25*).

**Figure 2:**
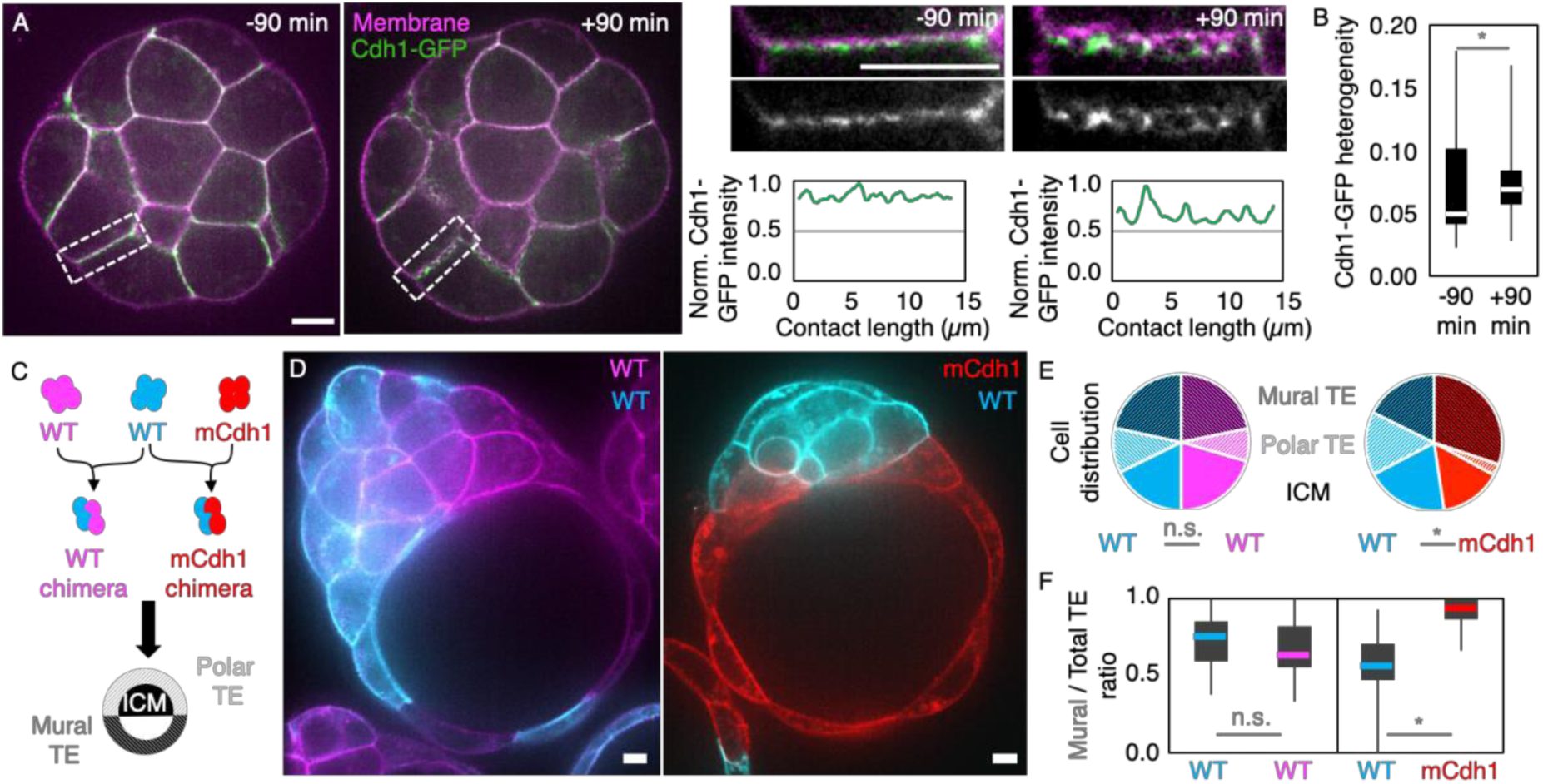
hydraulic fracture can be directed by cell adhesion. (A) Cdh1-GFP (green) and membrane (magenta) reorganize during microlumen formation. Left shows snapshots of the whole embryo 90 min before and 90 min after microlumen appearance. Right shows magnifications of the rectangles. Bottom right shows the intensity of Cdh1-GFP normalized to its maximal value along a 1 *µm* thick line following the cell-cell contact. Scale bars, 10 *µm.* (B) Coefficient of variation (standard deviation divided by the mean) of Cdh1-GFP intensity along cell-cell contacts. 228 and 221 contacts from 9 embryos at −90 and +90 min compared to the time of microlumen appearance respectively. * for Student’s t test at p < 10^−13^. (C) Schematic diagram of chimera experiments. (D) Chimeric embryos composed of control mTmG (cyan) and control mG (magenta) cells or of control mG (cyan) and mCdh1 (red) cells. Scale bars, 10 *µm.* (E-F) Distribution of cells in 22 control and 33 mutant chimeras. (E) Proportion of WT or mCdh1 cells in the mural (dark stripes) or polar (clear stripes) TE, or in the ICM (plain). Distribution heterogeneity between genotypes within each types of chimera is compared using Chi-squared test, * for p < 10^−2^, n. s. for not significant with p > 10^−2^. (F) Contribution to mural TE for each genotype. * for Mann-Whitney U test at p < 10^−3^, n. s. for not significant with p > 10^−2^.

Hydraulic fracture requires to build hydrostatic pressure between cells, first within microlumens and, eventually in the blastocoel. To evaluate the pressure in the blastocoel, we used micropipette aspiration, which allows measuring non-invasively the surface tension and hydrostatic pressure of liquid droplet-like objects (*26*). We measured pressures of 296 ± 114 Pa (Mean ± SD) for 25 blastocysts (Fig. S4), which is about 1O times higher than for individual blastomeres (*27, 28*). This reveals that the hydrostatic pressure in the blastocyst is large and of magnitude comparable to pressures capable of inducing hydraulic fracture in vitro (*25*). Therefore, fluid accumulation may be responsible for detaching cell-cell contacts and reorganizing Cdh1.

To prevent the accumulation of fluid coming from the outside medium at cell-cell contacts, we blocked the formation of the osmotic gradient that draws water inside the embryo, either by disrupting apicobasal polarity through Rock inhibition (*16*) or by directly increasing the osmolarity of the medium via the addition of sucrose (*29*). Either treatments prevented microlumen formation and Cdh1-GFP remained homogeneously distributed at cell-cell contacts (Fig. S5, Movies 3-4). This indicates that fluid accumulation is required for detaching cell-cell contacts and Cdh1 reorganization.

While fluid accumulation may displace Cdh1, conversely, Cdh1 could provide mechanical resistance to fluid accumulation. Therefore, we patterned cell-cell adhesion to test if this could affect lumen expansion. To achieve this, we took advantage of embryos in which Cdh1 is knocked out maternally (mCdh1), which form a blastocyst despite their initially lower adhesion (*27, 30*). We generated chimeric embryos made either from two differently labeled control embryos or from control and mCdh1 embryos (Fig. 2C). Control chimeras formed their blastocoel on either half of the embryo (Fig. 2D). On the other hand, in mCdh1 chimeras, the blastocoel formed preferentially on the mCdh1 half (Fig. 2D). This affected the allocation of cells to the different tissues of the blastocyst (Fig. 2E, Fig. S6) with mCdh1 blastomeres contributing almost exclusively to the mural part of the TE, which is the TE that lines the blastocoel (Fig. 2F). Therefore, patterning Cdh1 levels is sufficient to direct the accumulation of fluid. Altogether, we find that pressurized fluid collect along the path of lowest mechanical resistance, separates cell-cell contacts and reorganizes Cdh1 adhesion molecules. We therefore propose that microlumens form throughout the embryo by hydraulic fracture of cell-cell contacts.

We then proceeded to investigate how the embryo resolves the hundreds of microlumens into one single blastocoel during the discharge phase. One mechanism would be for microlumens to coalesce and fuse when in close proximity. On time scales ranging from tens of minutes to hours, we could not observe frequent fusion events or movements of microlumens towards the final position of the blastocoel (Movies 1-2). We used light-sheet microscopy to image microlumens at high temporal and spatial resolution, and observed that microlumens seem to drain their content on time scales of few minutes (Movie 5). Indeed, microlumens are physically connected through the intercellular space as revealed by labeled dextran diluted in the culture medium that builds up transiently at all cell-cell contacts during microlumen formation and then in the blastocoel (Movie 6). In fact, any difference in hydrostaitc pressure between two connected microlumens should lead to a flow of luminal fluid from the more pressurized microlumen to its counterpart (Fig. 3A). The pressure in microlumens is related to the tension and curvature at the lumen interface via the Young-Laplace equation (Supp. Text). At same microlumen tension, a pressure difference is due to differences in microlumen sizes. This leads smaller microlumens to discharge into larger ones (Fig. 3A, Movie 7), and may explain why small bicellular microlumens shrink earlier than larger multicellular microlumens (Fig. 1B, G). This process is reminiscent of Ostwald ripening (*31*), which drives the slow coarsening of foams or water-oil emulsions. In the embryo, however, the exchange is not limited by diffusion but rather by fluid flowing through the intercellular space. In addition, surface tension may not be homogeneous and, in fact, the direction of the flow may be reversed for a given asymmetry in tension between two lumens (Fig. 3B). To study microlumen dynamics, we modeled the embryo as a 2D network of pressurized microlumens and performed numerical simulations to predict statistically the position of the final lumen (Fig. 3C). Starting from random initial areas, the size of a microlumen evolves by direct swelling and through fluid exchange with its connected neighbors (Fig. 3C-D, Supp. Text). As observed experimentally (Fig. 1C), all simulations show a swelling phase followed by a coarsening of the network, which produces a single lumen that has siphoned all the fluid (Fig. 3D, Movie 8). Together, this supports a coarsening mechanism akin to Ostwald ripening (*31*).

**Fig. 3:**
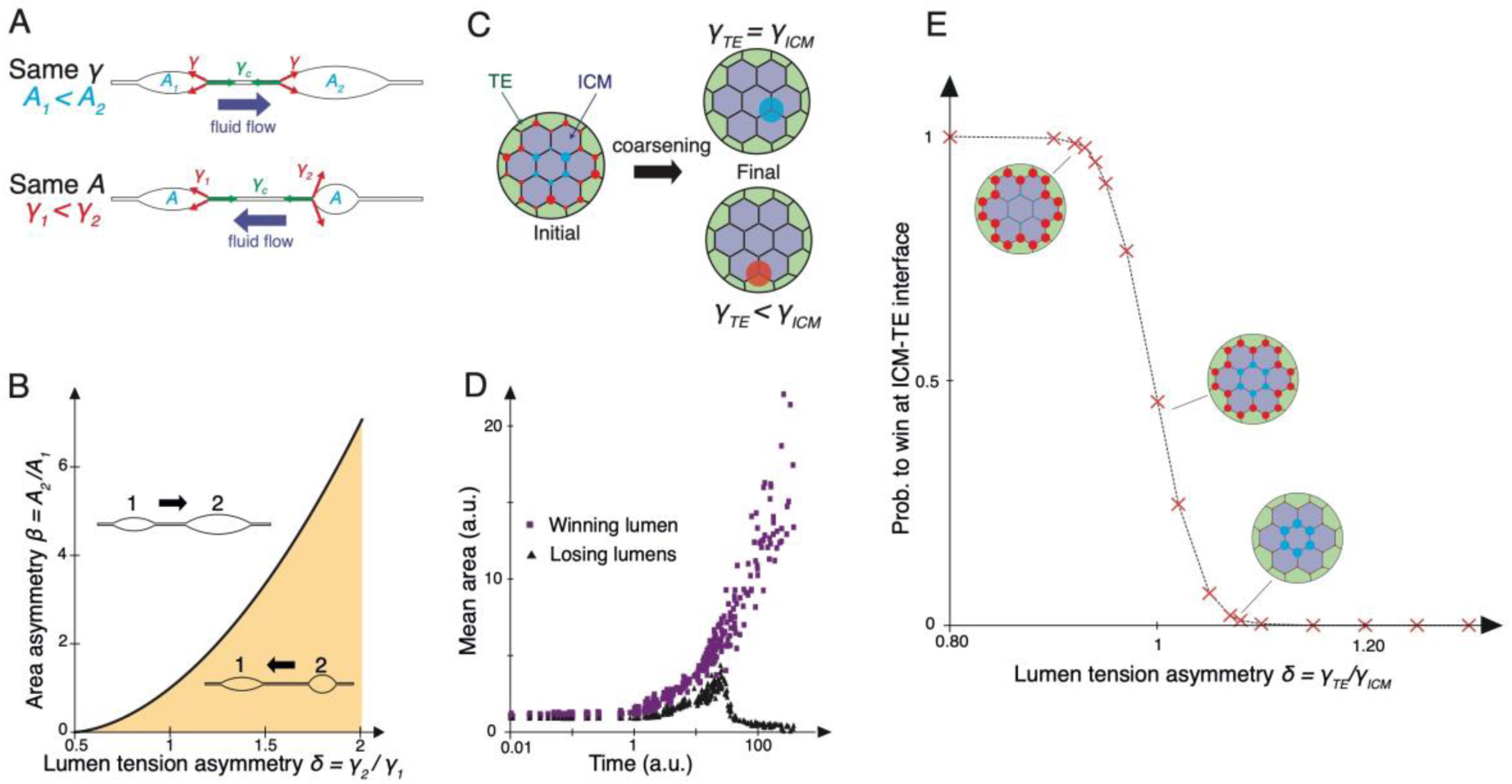
physical model of lumen coarsening and localization. (A) Schematic diagrams of two connected lumens in 2D. The fluid flow between the lumens is a function of their respective areas (***A***_*1*_, ***A***_*2*_) and lumen tensions (*γ _1_, γ _2_*). For equal tensions, the small lumen discharges into the larger one, while for equal areas, it is the lumen with higher tension that discharges. Adhesive contact tension *γ* _*c*_ is considered in Fig. S7. (B) Phase diagram for the fluid flow between two lumens as a function of their tension asγmmetrγ *δ=γ _2_* /*γ _1_* and initial size asγmmetrγ *β= A_2_ / A_1_.* Adhesive contact tension *γ*_*c*_ is 1. (C) Schematic diagram of a hexagonal network of lumens spanning the contacts between ICM and TE cells. Two lumen types either at ICM-ICM (blue) or ICM-TE interfaces (red) are associated with different tensions *γ*_*ICM*_ and *γ _TE_,* respectively. Lumen sizes are initially random (left) and the network coarsens until a single lumen, either at ICM-ICM or ICM-TE interface, has siphoned all the other lumens (right). (D) Time evolution of the mean area for winning (purple squares) and losing lumens (black triangles) from 20 representative simulations. (E) Winning probability for lumens at TE-ICM interfaces as a function of the tension asymmetry *δ = γ _TE_ / γ _ICM_.* Each data point results from averaging at least 5000 simulations on a regular hexagonal lattice. Insets show the mean localization of winning lumens.

In networks with homogeneous lumen tension, the lower connectivity of microlumens at the border favors the formation of the final lumen in the interior (Fig. 3E, Supp. Text). In agreement with mCdh1 chimera experiments (Fig. 2D), the model predicts that a pattern of adhesive contact tension could be sufficient to position the blastocoel at the TE-ICM interface (Fig. S7). Alternatively, when microlumens in the interior are imposed a small excess in tension, the final lumen ends up invariably at the margin (Fig. 3E). The model predicts hence that higher lumen tension at the ICM-ICM interface is sufficient to position the blastocoel between TE and ICM cells, as observed in vivo.

To examine whether differences in surface tension between blastomeres could explain how ripening directs luminal fluid towards the TE-ICM interface, we first investigated the shape of microlumens. When measuring the radius of curvature of microlumens, we detect asymmetries, which are more pronounced at heterotypic TE-ICM interfaces compared to those at homotypic TE-TE interfaces (Mean ± SEM, curvature ratio of 1.01 ± 0.02 for 71 TE-TE microlumens and 1.25 ± 0.04 for 58 TE-ICM microlumens from 7 embryos, Student t test p < 10^−2^, Fig. 4A-B, Movie 1). Moreover, microlumen at TE-ICM interfaces bulge into the TE cell in most cases rather than into ICM cells (79% of 58 microlumens from 7 embryos, Chi squared p < 10^−3^)This suggests differences in the hydrostatic pressure and/or surface tension between TE and ICM cells. Such differences are supported by the cell-scale curvature of TE-ICM interfaces, where ICM cells bulge into TE cells (Fig. **S8).** This disparity is likely due to higher contractility of ICM cells compared to TE cells, which is responsible for their sorting at the 16-cell stage (*28*). Indeed, inhibiting cell contractility decreases the curvature at TE-ICM interfaces (Fig. S8) and the surface tension of the blastocyst (Fig. S4). Our model predicts that this pattern of contractility within the embryo would result in microlumens preferentially discharging to the TE-ICM interface (Fig. 3E, Movie 8).

**Figure 4:**
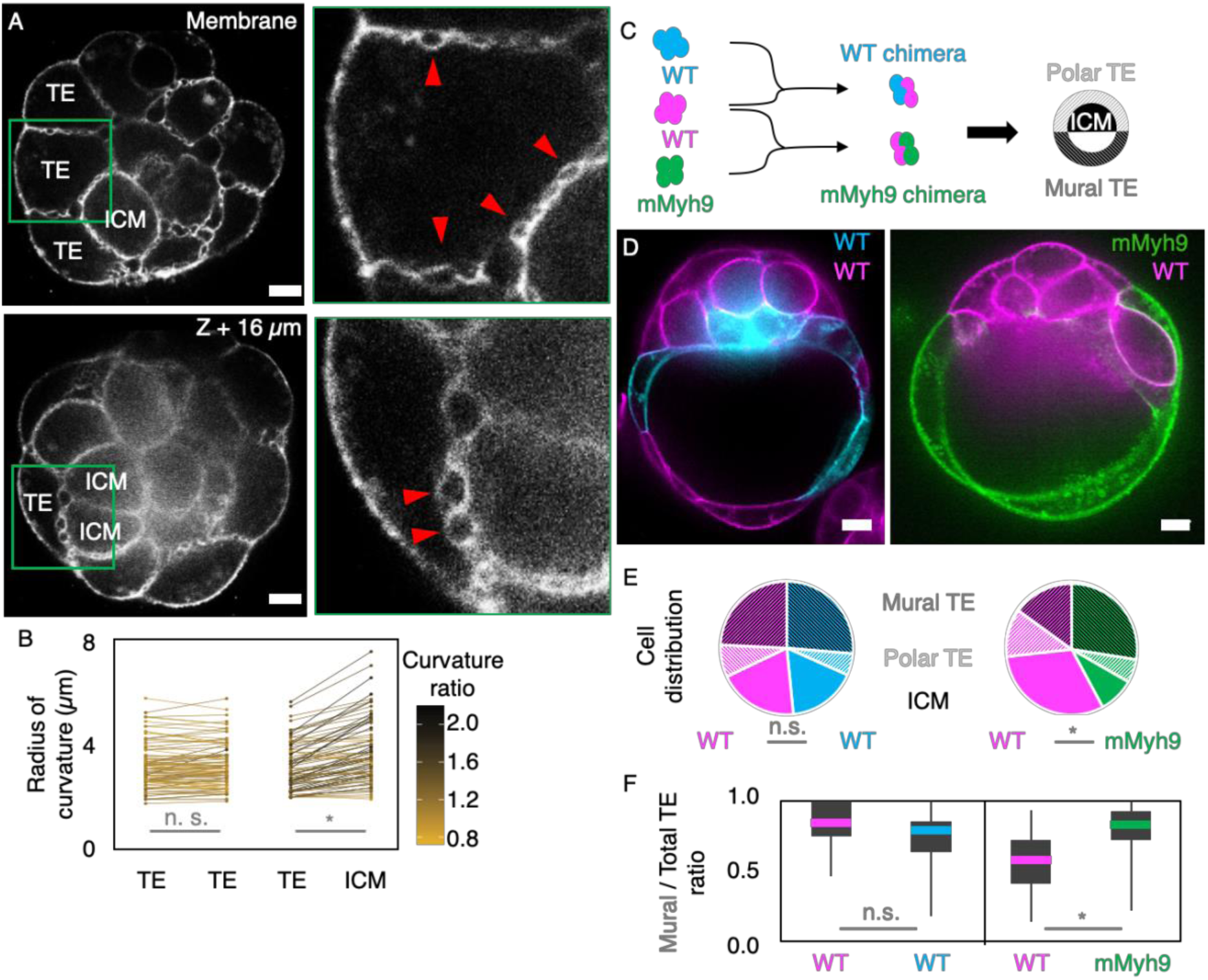
microlumen coarsening is controlled by cell contractility. (A) The curvature of microlumens is more symmetric at TE-TE interfaces (top, arrowheads) and more asymmetric at TE-ICM interfaces (bottom image shows a confocal slice 16 *µ*m below the one above). Green squares are magnified 3 times on the right images. Scale bar, 10 *µ*m. (B) Radius of curvature at microlumens facing TE-TE (71) and TE-ICM (58) interfaces from 7 embryos during the discharge phase, 60 minutes after inversion. The ratio of curvature is color-coded with TE cells ordered randomly for TE-TE ratios. * for paired Student’s *t* test at p < 10^−7^, n. s. for not significant with p > 10^−2^. (C) Schematic diagram of chimera experiments. (E-F) Distribution of cells in 25 control and 28 mutant chimeras. (E) Proportion of WT or mMyh9 cells in the mural (dark stripes) or polar (clear stripes) TE, or in the ICM (plain). Distribution heterogeneity between genotypes within each types of chimera is compared using Chi-squared test, * for p < 10^−2^, n. s. for not significant with p > 10^−2^. (F) Contribution to mural TE for each genotype. * for Mann-Whitney U test at p < 10^−3^, n. s. for not significant with p > 10^−2^.

To experimentally test the ability for heterogeneous contractility to direct the offloading of microlumens, we took advantage of embryos in which Myh9 is maternally knocked out (mMyh9), which are viable despite their lower levels of Myh9 and lower surface tension (*28*). We generated chimeric embryos made either from two differently labeled control embryos or with mMyh9 embryos in order to pattern cell contractility (Fig. 4C). As previously (Fig. 2C-F), control chimeras form their blastocoel on either side of the embryo (Fig. 4D). On the other hand, in mMyh9 chimeras, the blastocoel forms preferentially on the mMyh9 half (Fig. 4D). This affects the allocation of cells to the different tissues of the blastocyst (Fig. 4E). In agreement with previous analogous experiments (*28*), mMyh9 blastomeres are depleted from the ICM (Fig. S6) and, instead, contribute mostly to the mural TE (Fig. 4F). Therefore, differences in Myh9 levels are sufficient to control the position of the blastocoel, in agreement with corresponding simulations (Movie 9). Altogether, we propose that a mechanism akin to Ostwald ripening directs luminal fluid to unload microlumens into the blastocoel.

We have established that both patterned cell-cell adhesion and patterned cell contractility can position the blastocoel (Fig. 2D-F, 4D-F). This suggests that both localized tracking of cell-cell contacts and directed fluid offloading could set the first axis of symmetry of the mouse embryo. These two phenomena constitute complementary mechanisms explaining the formation and positioning of lumens in general, and of basolateral lumens in particular, which can be found in healthy (*32, 33*) and pathological situations (*15*). As far as the blastocyst is concerned, although both tracking and ripening can position the blastocoel, we are unable to detect any conclusive relation between the final position of the blastocoel and the initial localization of Cdh1 or contact fracture (Fig. S2-3, S9). On the other hand, differences in contractility are undoubtedly at play (*28, 34,35*) (Fig. 4, Fig. S8). Therefore, we propose that the first axis of symmetry of the mouse embryo is positioned by contractility-mediated ripening of microlumens formed after tracking of cell-cell contacts.

Determining the molecular and mechanical events controlling the formation of microlumens and the fracture of cell-cell contacts constitutes exciting new research avenues, which will greatly benefit from previous studies on contact mechanical stability (*25, 36*). Finally, to build a comprehensive model of blastocoel formation, future studies will need to integrate explicitly individual cell mechanics1 and to evaluate the contributions of other cellular processes such as vesicular trafficking (*18, 37*) and ion exchange (*10*).

---

**Supplementary Figure 1:**
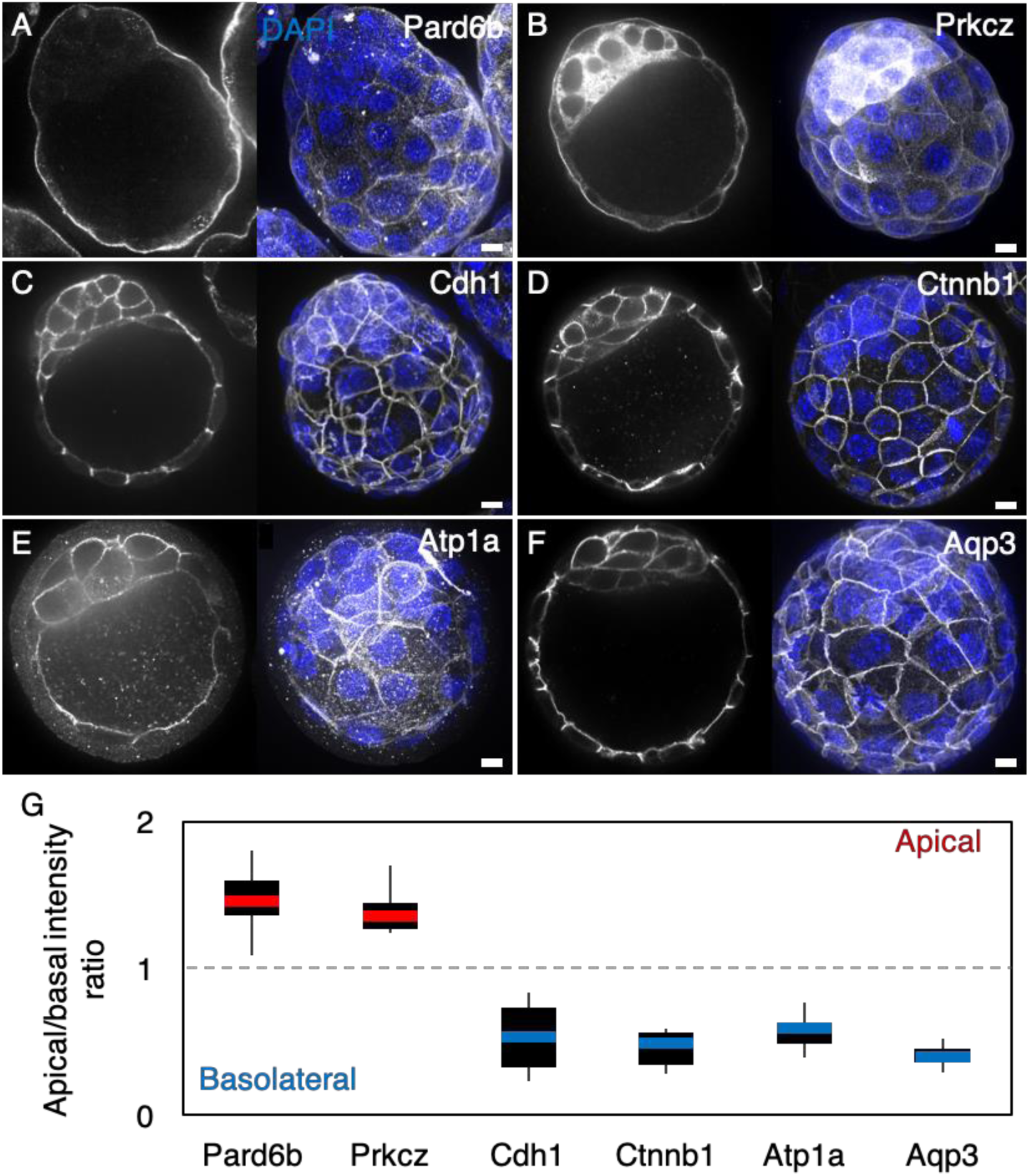
blastocyst form the blastocoel on the basolateral interface of cells. (A-D) Immunostaining (grey) showing single confocal slices (left) and maximum projections (right) with nuclei stained with DAPI (blue). (A-B) Apical markers Pard6b and Prkcz. (C-D) Basolateral markers Cdh1, Ctnnb1, Atp1a, Aqp3. (G) Apical to basal intensity ratio for 3 mural TE cells from 15/14/10/11/8/16 embryos for Pard6b, Prkcz, Cdh1, Ctnnb1, Atp1a, Aqp3 respectively.

**Supplementary Figure 2:**
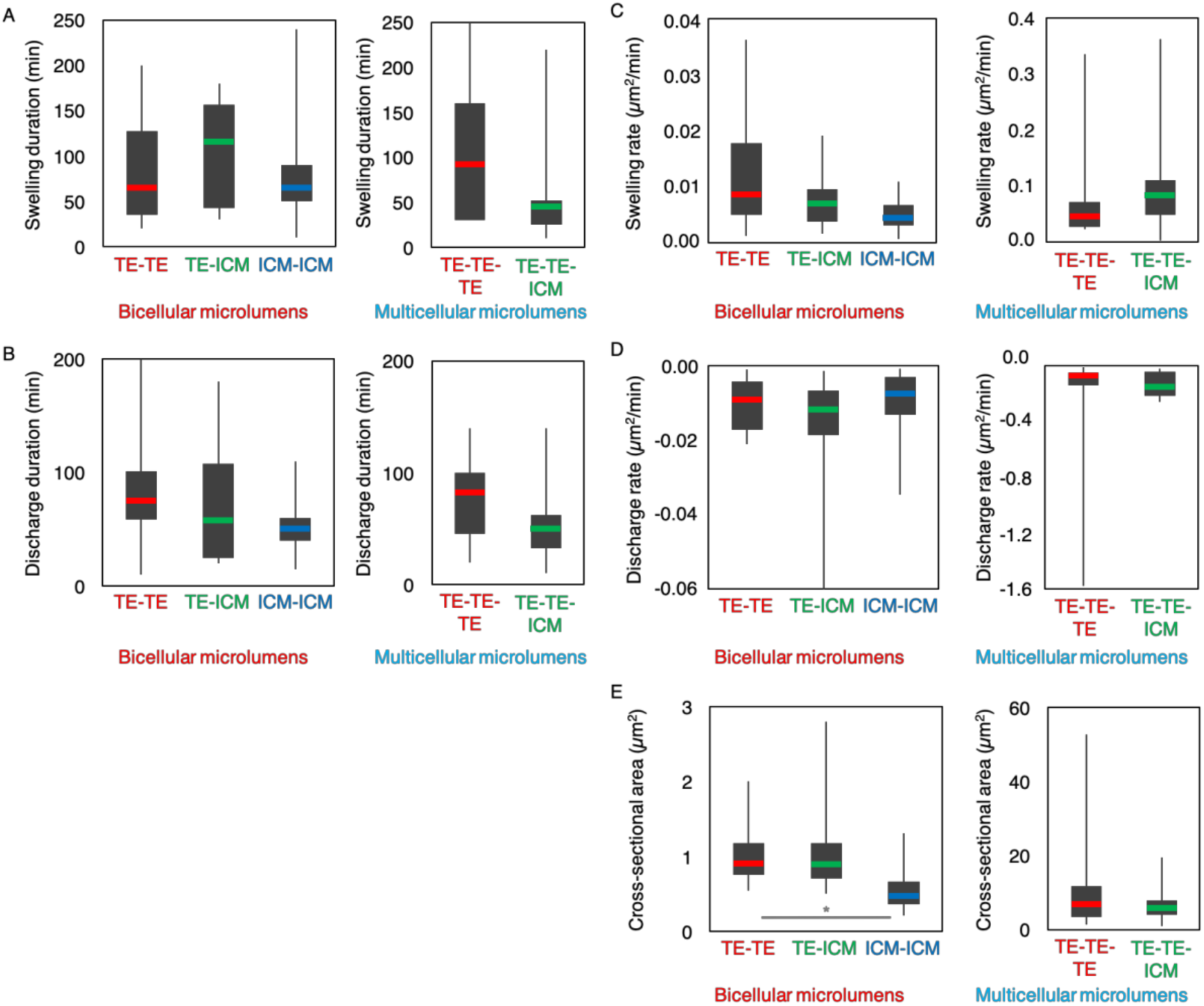
microlumen dynamics are homogeneous at different interfaces. (A-E) Swelling (A) and discharge duration (B), swelling (C) and discharge rate (D), and cross-sectional area (E) for bicellular microlumens at 12 TE-TE, 10 TE-ICM, 13 ICM-ICM interfaces and for multicellular microlumens at 8 TE-TE-TE and 11 TE-TE-ICM junctions from 7 embryos. Student’s t test at p > 10^−2^ gives no significant differences.

**Supplementary Figure 3:**
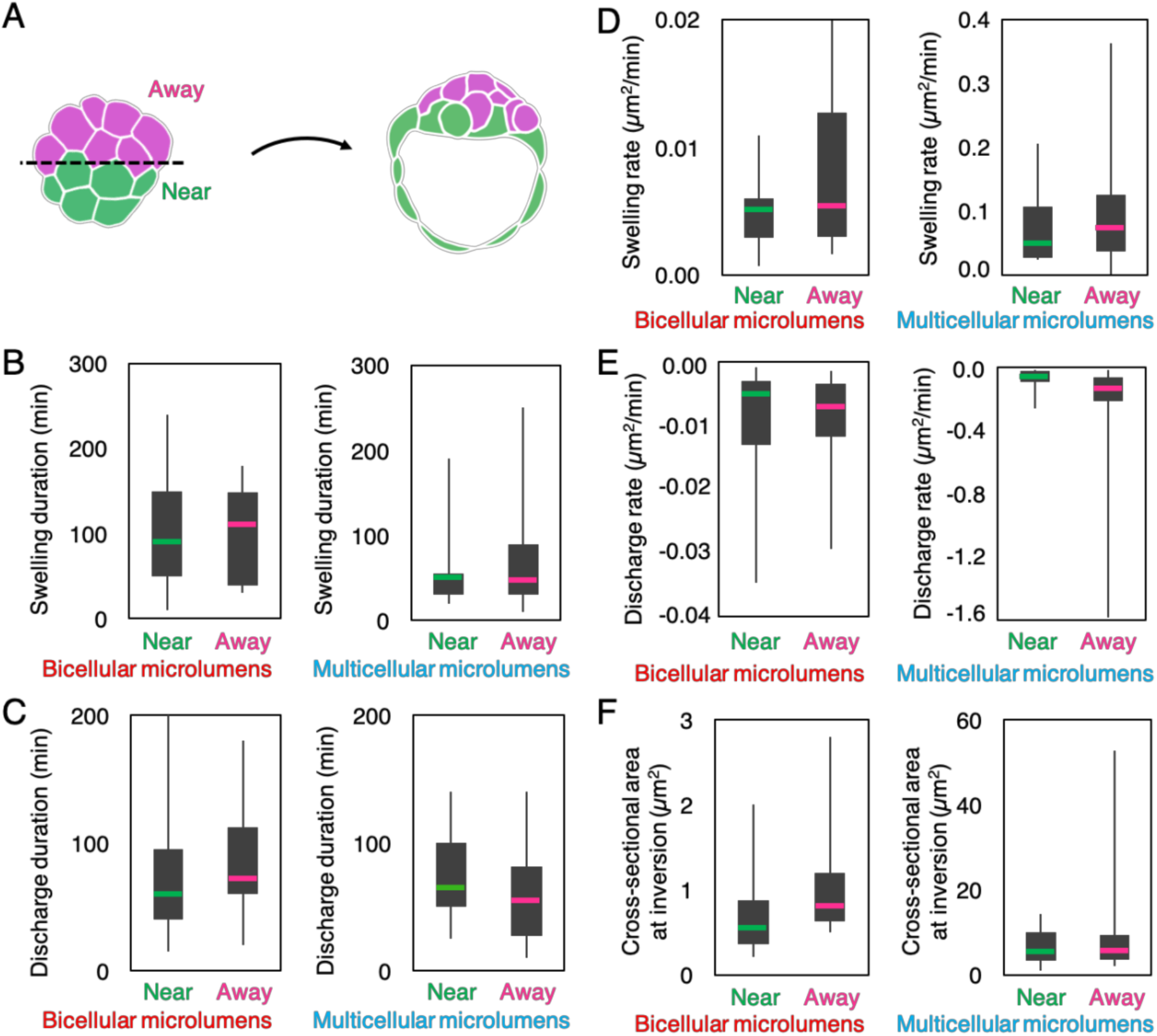
microlumen dynamics are homogeneous throughout the embryo. (A) Schematic diagram of the location of the measurements of microlumen dynamics. (B-F) Swelling (B) and discharge duration (C), swelling (D) and discharge rate (E), and microlumen cross-sectional area (F) for interfaces near (N = 13) or away (N = 12) from the final position of the blastocoel in 5 embryos for bicellular microlumens and for interfaces near (N = 9) or away (N = 12) from the final position of the blastocoel in 7 embryos for multicellular microlumens. Student’s t test at p > 10^−2^ gives no significant differences.

**Supplementary Figure 4:**
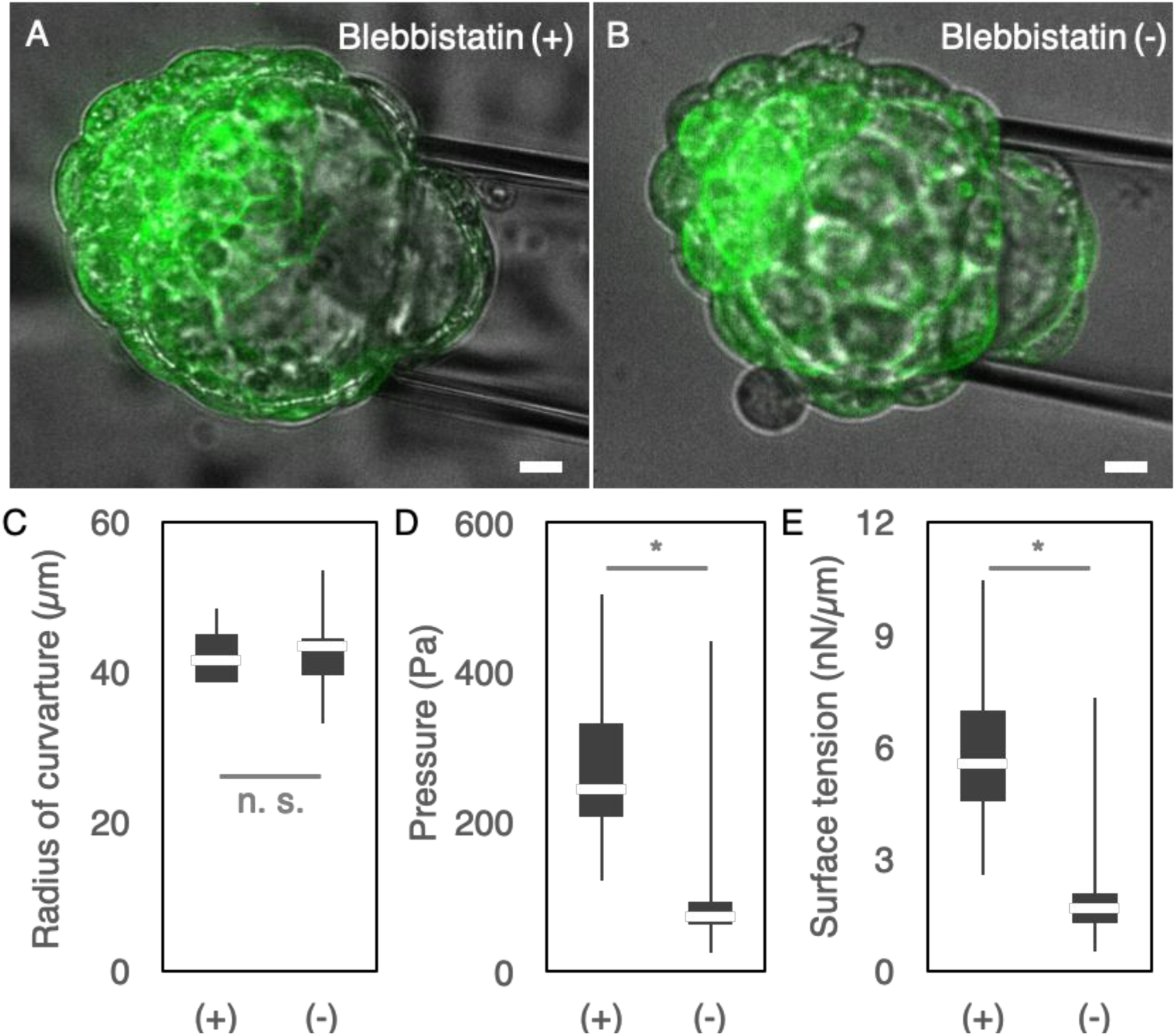
hydrostatic pressure and surface tension of the blastocyst are controlled by actomyosin contractility. (A-B) Blastocyst microaspiration on the mural TE in the presence of 25 µM inactive enantiomere Blebbistatin (+) or active enantiomere Blebbistatin (-). Brighfield in grey and membrane (mTmG) in green. Scale bar, 10 *µ*m. (C-E) Radius of curvature of the mural TE, hydrostatic pressure and surface tension of 25 (+) and 24 (-) blastocysts. * for Student’s t test at p < 10^−6^, n. s. for not significant with p > 10^−2^.

**Supplementary Figure 5:**
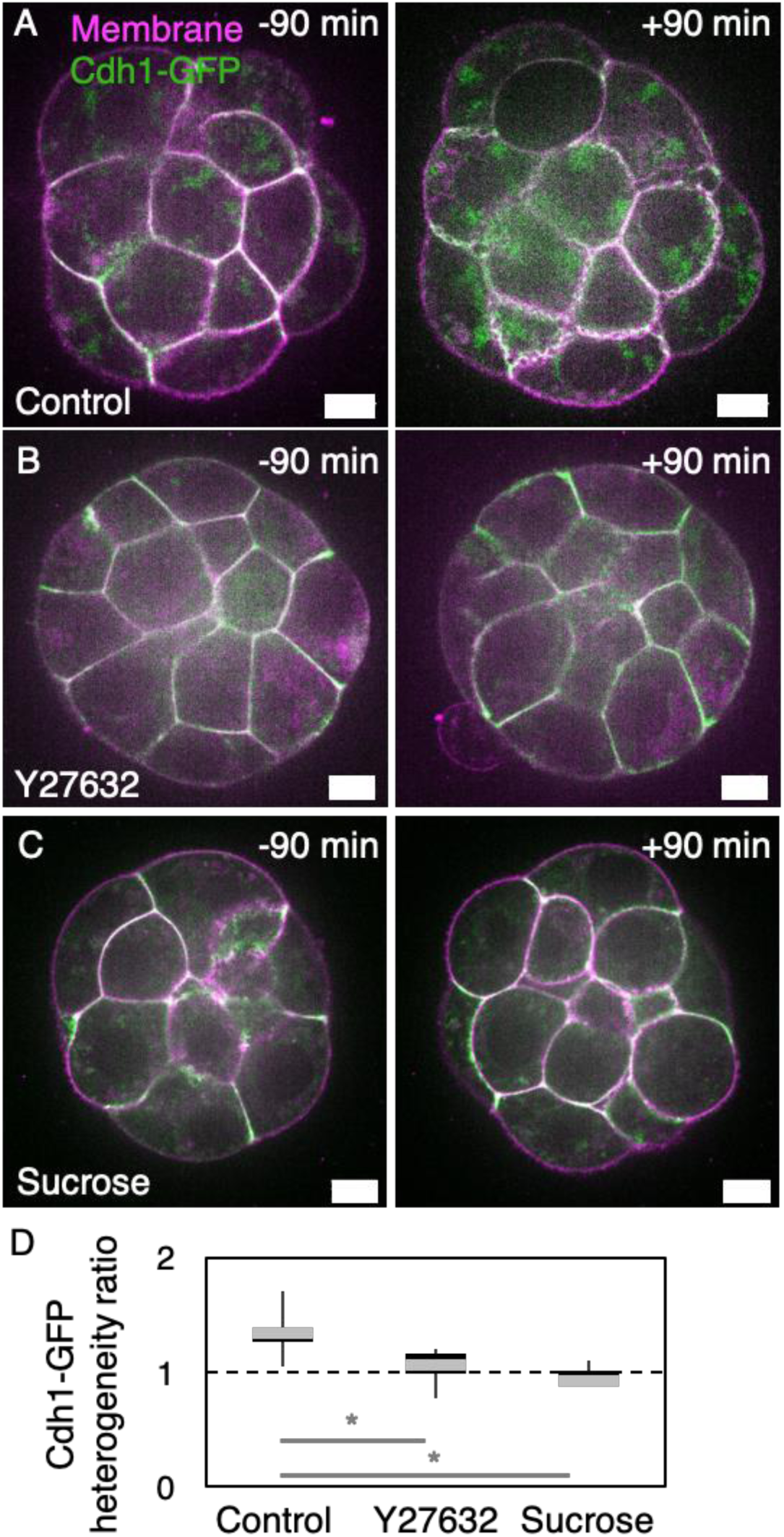
fluid accumulation is required for Cdh1-GFP rearrangement. (A-C) Cdh1-GFP (green) and membrane (magenta) for embryos in control medium (A), 20 µM Y27632 (B) or 175 mM sucrose (C) containing medium. Scale bar, 10 *µ*m. (D) Quantification of Cdh1-GFP signal reorganisation along cell-cell contacts. The ratio of coefficient of variation is calculated between 90 min before and 90 min after the appearance of microlumens in control embryos and at the time of the latest control embryo from the same experiment in Y27632 and sucrose treated embryos. 449 *I* 298 *I* 186 contacts from 13 / 10 / 3 embryos for control, Y27632 and sucrose respectively. * for Student’s t test at p < 10^−2^.

**Supplementary Figure 6:**
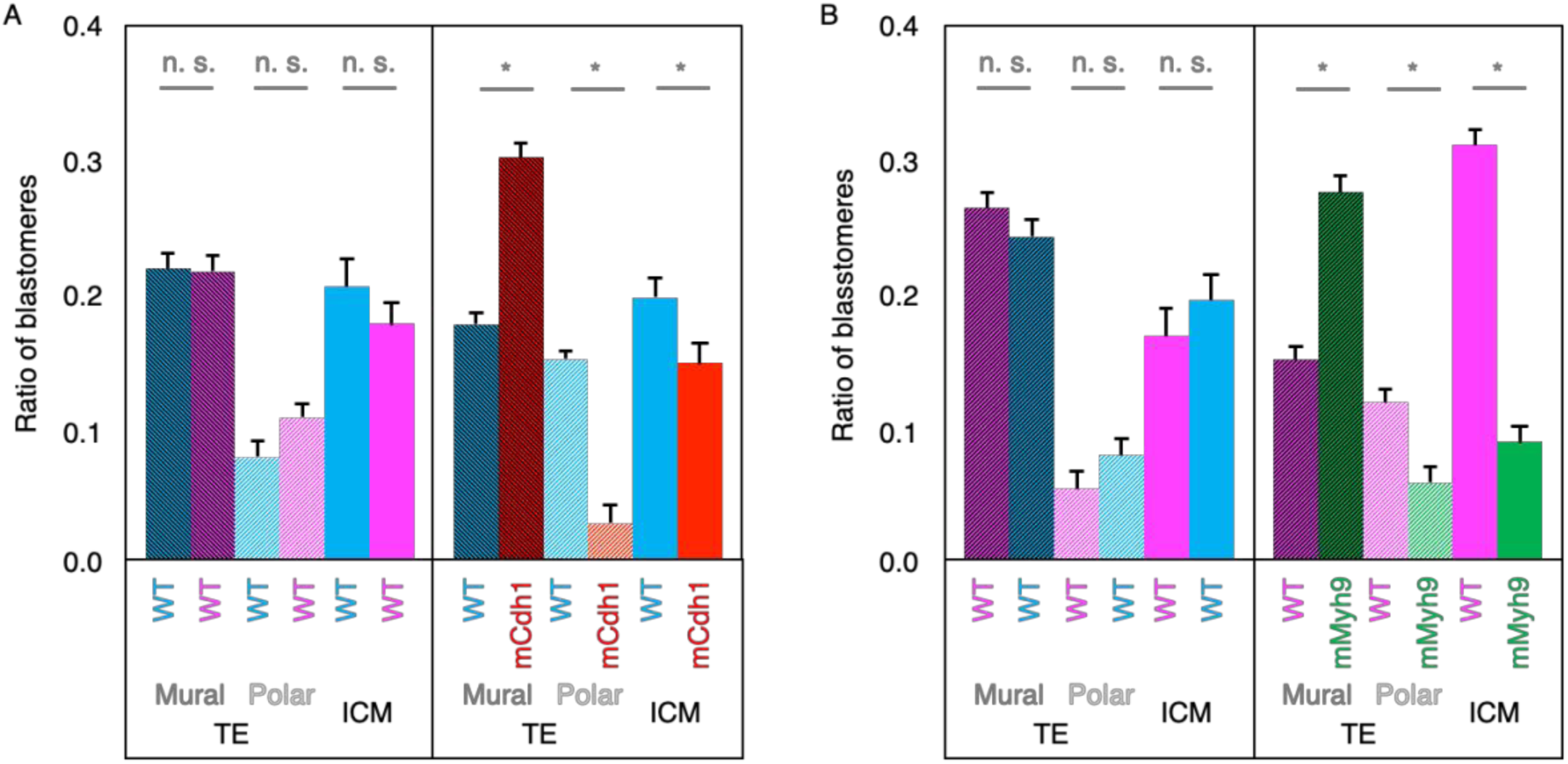
cell distribution in chimeric embryos. (A-B) Mean± SEM contribution of each genotype to mural (dark stripes) or polar (clear stripes) TE, or to ICM (plain). (A) Distribution of cells in 22 control and 33 mCdh1 mutant chimeras. (B) Distribution of cells in 25 control and 28 mMyh9 mutant chimeras. * for Mann-Whitney U test at p < 10^−3^, n.s. for not significant at p > 10 ^−2^

**Supplementary Figure 7:**
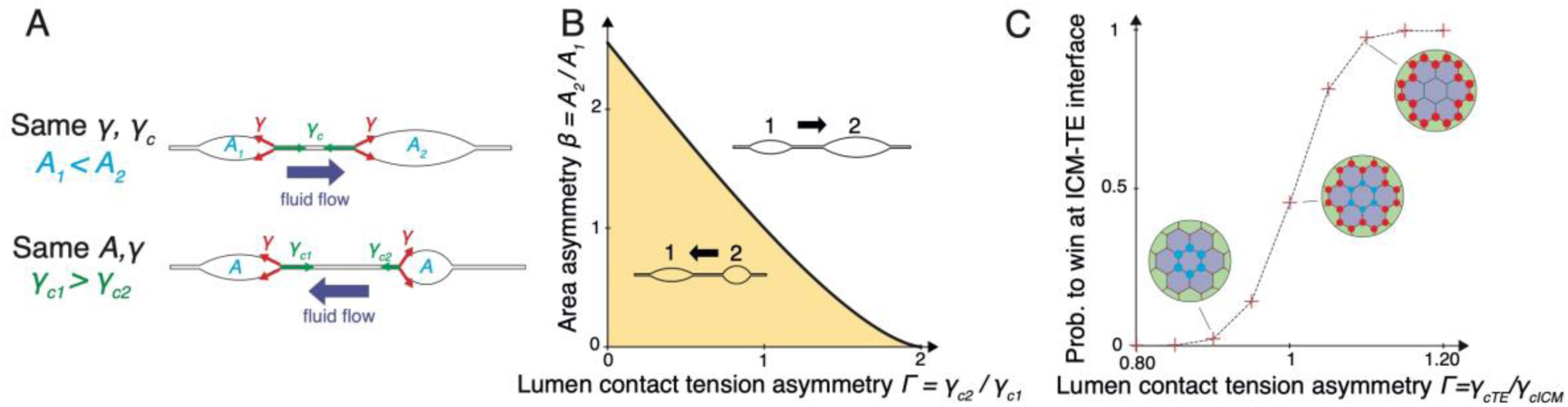
effect of adhesive contact tension on lumen coarsening and localization. (A) Schematic diagrams of the evolution of two connected lumens in 2D. The fluid flow between lumens is a function of their respective areas (***A***_*1*_, ***A***_*2*_) and adhesive contact tensions (*γ*_*c1*_*,γ*_*c2*_). For equal contact tensions, the small lumen discharges into the larger one, while for equal areas, it is the lumen for which the contact has a lower tension that discharges. *γ* is the tension at the lumen. (B) Phase diagram for the fluid flow between two lumens as function of the adhesive contact tension asymmetry *Γ= γ* _*c2*_ */ γ* _*c1*_ and initial size asymmetry *β =* ***A***_*2*_*/* ***A***_*1*_. Lumen tension γ is 1. (C) Winning probability for lumens at TE-ICM interfaces as a function of the adhesive contact tension asymmetry *Γ = γ* _*cTE*_ */γ*_*cicM*_ Each data point results from averaging at least 5000 simulations on a regular hexagonal lattice. Insets show the mean localization of winning lumens.

**Supplementary Figure 8:**
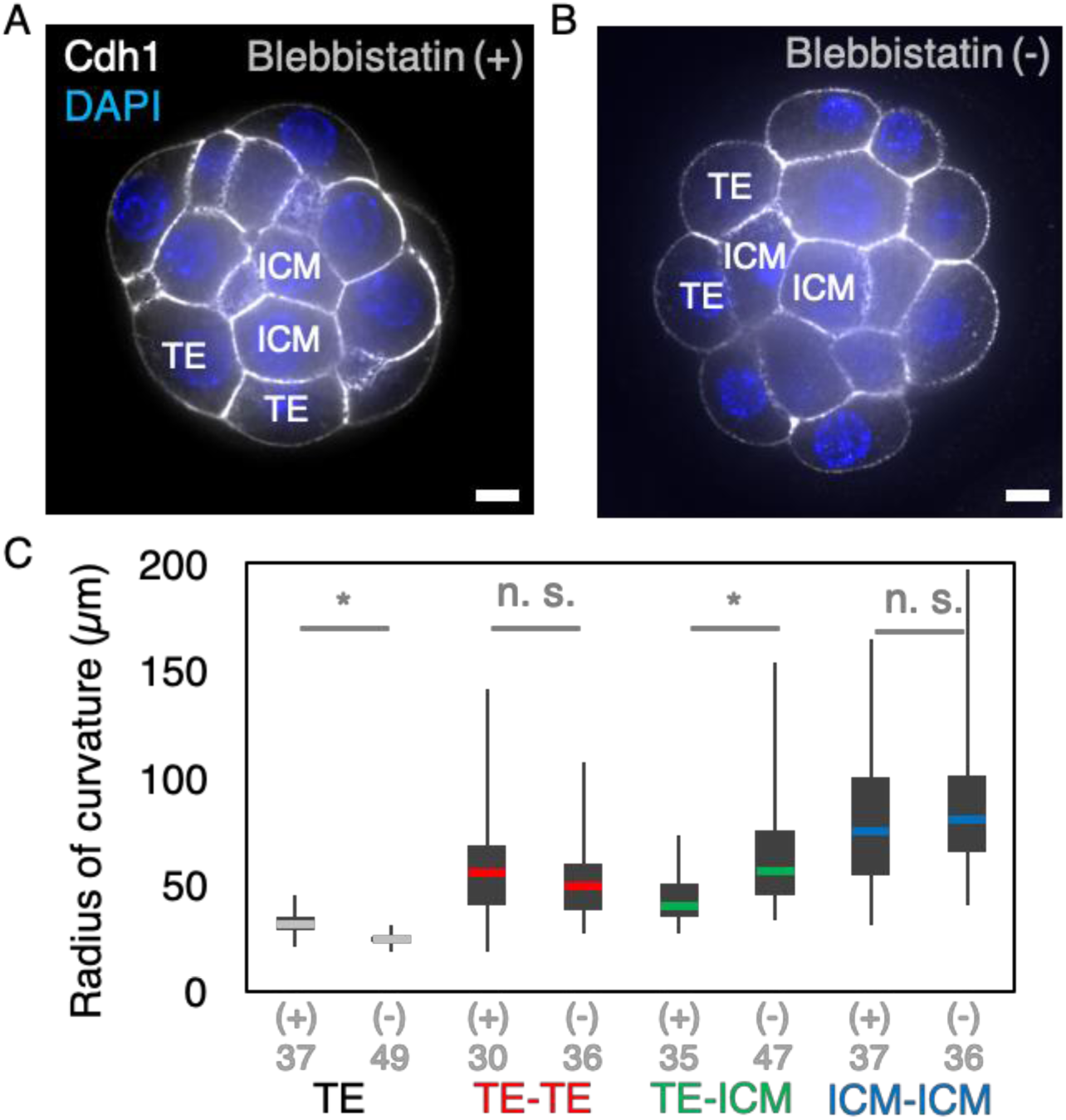
different contractility of TE and ICM cells curve TE-ICM interfaces. (A-B) 32-cell stage embryos before lumen formation in 25 *µ*M inactive enantiomere Blebbistatin (+) or active enantiomere Blebbistatin (-). Scale bar, 10 *µ*m. (C) Radius of curvature at TE-TE, TE-ICM and ICM-ICM interfaces, and TE interfaces with the medium of 8 Blebbistatin (+) and 9 Blebbistatin (-) treated embryos. The number of interfaces is indicated. * for Student’s t test at p < 10^−3^, n. s. for not significant with p > 10^−2^.

**Supplementary Figure 9:**
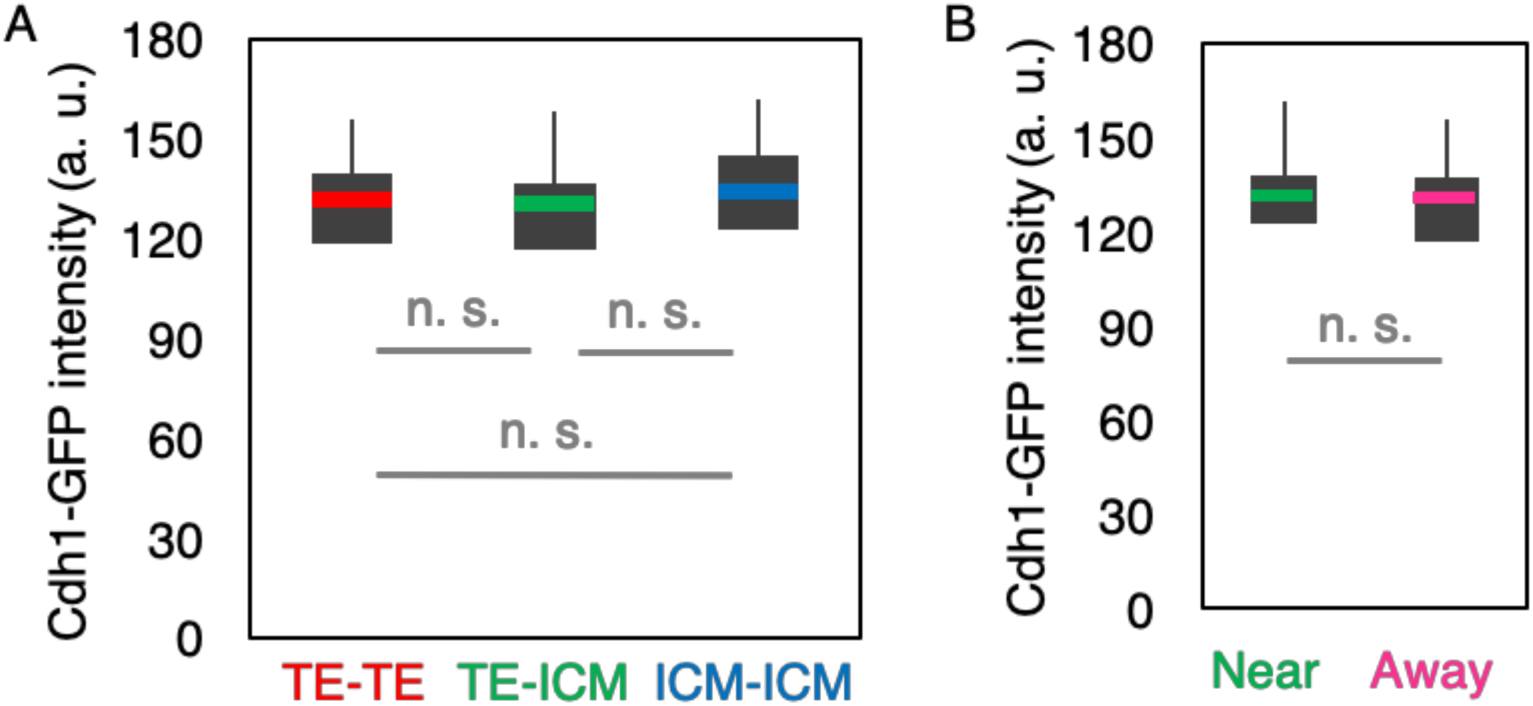
adhesion is homogeneous throughout the embryo. Mean Cdh1-GFP intensity at 21 TE-TE, 34 TE-ICM and 8 ICM-ICM interfaces from 10 embryos. Student’s t test at p > 10^−2^ gives no significant differences. (B) Mean Cdh1-GFP intensity at cell-cell contacts near (N = 33) or away (N = 30) from the final position of the blastocoel in 10 embryos. Student’s t test at p > 10^−2^ gives no significant differences.

**Movie 1: swelling and shrinkage of microlumens during blastocoel formation.**

Single confocal slice of a timelapse of blastocoel formation in a mTmG embryo showing plasma membrane labelling. Images taken every 10 minutes, scale bar 10 *µ*m. Time is set relative to the time of appearance of the first microlumens.

**Movie 2: Cdh1 reorganization during microlumen formation.**

Timelapse of blastocoel formation in a Cdh1-GFP; mTmG embryo showing Cdh1 (green) and plasma membrane (magenta) labelling on two confocal slices separated by 20 *µ*m. Images taken every 30 minutes, scale bar 10 *µ*m.

**Movie 3: Rock inhibition prevents microlumen formation and Cdh1 reorganization.**

Single confocal slice of a timelapse of blastocoel formation in a Cdh1-GFP; mTmG embryo showing Cdh1 (green) and plasma membrane (magenta) labelling in the presence of 20 *µ*M Y27632. Images taken every 30 minutes, scale bar 10 *µ*m.

**Movie 4: hyperosmosis prevents microlumen formation and Cdh1 reorganization.**

Single confocal slice of a timelapse of blastocoel formation in a Cdh1-GFP; mTmG embryo showing Cdh1 (green) and plasma membrane (magenta) labelling in the presence of 175 mM sucrose. Images taken every 30 minutes, scale bar 10 *µ*m.

**Movie 5: high resolution imaging of the discharge of microlumens.**

Projection of confocal slices of a timelapse of blastocoel formation in a mTmG embryo showing plasma membrane labelling. The central microlumen expands as microlumens from neighbouring cell-cell interfaces shrinkage. Images taken every 5 s, scale bar 10 *µ*m.

**Movie 6: high resolution imaging of fluid accumulation.**

Single confocal slices of a timelapse of blastocoel formation in a mTmG embryo showing plasma membrane labelling (cyan) growing in a medium containing a 3 kDa Dextran coupled to Alexa 488 (red). Labelled dextran accumulates transiently at all cell-cell contacts during microlumen formation and then in the blastocoel. Accumulation of labelled dextran at contacts between ICM cells indicate that intercellular space is connected. Images taken every 2 min, scale bar **1**0 *µ*m.

**Movie 7: simulation of two lumens exchanging fluid through a pipe.**

Both simulations have the same area asymmetry (A_2_ = 2A_1_) but lumen tension is either symmetric γ _1_ = γ _2_ = **1** (top), or asymmetric 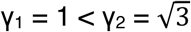 (bottom). Contact tension *γ*_*c*_= 1

**Movie 8: simulation of coarsening in a regular hexagonal network of lumens similar to Fig. 3C.**

Different tensions are imposed at ICM-ICM (blue, γ _TE_ = 0. 9) and ICM-TE (red, γ _ICM_ = **1)** interfaces. Contact tension γ_c_= **1.**

**Movie 9: simulation of the coarsening in a network of lumen representing a chimeric embryo with low contractility mutant cells.**

Mutant TE cells (light green) have a lower contractility than WT TE cells (light magenta): lumen tensions are γ _mutant_ = 0. 7 at the mutant/ICM (green), γ _TE-mutant_ = [γ _mutant_+γ_TE_]/2 at the TE/mutant/ICM interface (grey), γ _TE_ = 0. 9 at TE/ICM interface (magenta) and γ _ICM_ = 1 at the ICM interface (blue). Contact tension *γ*_*c*_= 1

## Acknowledgements

We are grateful to the imaging platform of the Genetics and Developmental Biology unit at lnstitut Curie (PICT-IBiSA@BDD) for their outstanding support. We thank the animal facility of Institut Curie for their invaluable help.

We are grateful to the Buchholz laboratory for providing the Cdh1-GFP BAC construct. We thank Yvonne Petersen from the transgenesis platform at the European Molecular Biology Laboratory, Takashi Hiiragi and his lab for their very generous help in making the Cdh1-GFP mouse line.

We thank the Maître and Turlier labs, Yohanns Bellaïche, Andrew Clark and Guillaume Charras for fruitful discussions and comments on the manuscript.

Research in the lab of JLM, who is supported by the Institut Curie, the CNRS and the INSEAM, is funded by the ATIP-Avenir program, an ERC-2017-StG 757557, a PSL “nouvelle équipe” grant and Labex DEEP (ANR-11-LBX-0044) which are part of the IDEX PSL (ANR-10-IDEX-0001-02 PSL). The lab of HT acknowledges support from the Fondation Bettencourt-Schueller, the CNRS-INSERM ATIP-Avenir program and the Collège de France.

## Authors contributions

HT and JLM conceptualized the project and acquired funding. JGD and JLM designed experiments. JGD, AFT and LOP performed experiments. JGD, AFT, LOP and JLM analysed the data. MLVS, AM and HT designed the theoretical model and performed the numerical simulations.

## Methods

### Embryo work

#### Recovery and culture

All animal work is performed in the animal facility at the Institut Curie, with permission by the institutional veterinarian overseeing the operation (APAFIS #11054-2017082914226001). The animal facilities are operated according to international animal welfare rules.

Embryos are isolated from superovulated female mice mated with male mice. Superovulation of female mice is induced by intraperitoneal injection of 5 international units (IU) pregnant mare’s serum gonadotropin (PMSG, Ceva, Syncro-part), followed by intraperitoneal injection of 5 IU human chorionic gonadotropin (hCG, MSD Animal Health, Chorulon) 44-48 hours later. Embryos are recovered at E1.5 or E2.5 by flushing oviducts from plugged females with 37°C FHM (LifeGlobal, ZEHP-050 or Millipore, MR-122-D) using a modified syringe (Acufirm, 1400 LL 23).

Embryos are handled using an aspirator tube (Sigma, A5177-5EA) equipped with a glass pipette pulled from glass micropipettes (Blaubrand intraMark or Warner Instruments).

Embryos are placed in KSOM (LifeGlobal, ZEKS-050 or Millipore, MR-107-D) or FHM supplemented with 0.1 % BSA (Sigma, A3311) in 10 *µ*L droplets covered in mineral oil (Sigma, M8410 or Acros Organics). Embryos are cultured in an incubator with a humidified atmosphere supplemented with 5% CO_2_ at 37°C.

To remove the Zona Pellucida (ZP), embryos are incubated for 45-60 s in pronase (Sigma, P8811).

For imaging, embryos are placed in 5 or 10 cm glass-bottom dishes (MatTek).

#### Mouse lines

Mice are used from 5 weeks old on.

(C57BL/6xC3H) F1 hybrid strain is used for wild-type (WT).

To visualize plasma membranes, mTmG or mG (Gt(ROSA)26Sortm4^(ACTB-tdTomato,-EGFP)Luo^) mice are used (*22*). To remove LoxP sites specifically in oocytes, Zp3-cre (Tg(Zp3-cre)93Knw) mice are used (*38*). To generate mCdh1 embryos, Cdh1^tm^2^kem^ mice are used (*39*) to breed Cdh1^tm^2^kem/tm^2^kem^; z p3^Cre/+^ mothers with mTmG or mG fathers. To generate mMyh9 embryos, Myh9^tm^5^RSad^ mice are used (*40*) to breed ^tm^5^Rsad/tm^5^RSad^; Zp3^cre/+^ mothers with mTmG or mG fathers.

To visualize Cdh1, transgenic Cdh1-GFP mice were generated by the micro-injection of a bacterial artificial chromosome containing the Cdh1 gene modified by recombineering (RP23-262N14) (*24, 41*) into (CD1xC57BL/6) F1 hybrid zygotes that were transferred into pseudo-pregnant CD1 female mice. Founder mice were examined for the presence of BAC integration by genotyping using CACATGAAGCAGCACGACTT and AGTTCACCTTGATGCCGTTC primers amplifying 250bp of the BAC backbone and GACCACA TGAAGCAGCACGAC and CGAACTCCAGCAGGACCATG primers amplifying 445bp of the GFP tag. Preimplantation embryos from positive mice were imaged to verify the junctional localization of the GFP signal in live embryos and in embryos stained using Cdh1 antibody. Mice were outcrossed to (C57BL/6xC3H) F1 hybrid mice to generate N1.

#### Chemical reagents

Blebbistatin (+), an inactive enantiomere of the inhibitor, or (-), the selective inhibitor myosin II ATPase activity, (Tocris, 1853 and 1852) 50 mM DMSO stocks are diluted to 25 *µ*M in KSOM or FHM. 32-cell stage embryos are incubated in medium containing Blebbistatin without covering with mineral oil for 1 h before the beginning of the experiment.

Y27632, a selective inhibitor of ROCK (Tocris, 1254), 100 mM PBS stock is diluted to 10 *µ*M in KSOM. Late 16-cell stage and early 32-cell stage embryos are incubated and imaged overnight.

KSOM containing 175 mM sucrose is made by diluting a 1.4 M sucrose KSOM preparation adapted from (*42*) into commercial KSOM. Late 16-cell stage and early 32-cell stage embryos are incubated and imaged overnight.

Alexa Fluor 488 coupled to a 3 kDa Dextran (Sigma, D34682) is added to KSOM at 0.1 g.L^−1^ Embryos are placed in labelled Dextran at the 16-cell stage before tight junctions fully seal.

#### Chimeric embryos

To build chimeras, 2- and 4-cell stage embryos are removed from their ZP using pronase and placed into Ca^2+^-free KSOM for 2-3 min before being aspirated multiple times through a glass pipette until dissociation of cells into singlets or doublets. Using a mouth pipette, one 2-cell stage or two 4-cell stage blastomeres from two different embryos are then re-aggregated to form a complete embryo. Only embryos developing to blastocyst stage are considered and we exclude chimeras in which fewer than half of the cells come from one of the two original embryos.

#### lmmunostaining

Embryos are fixed in 2% PFA (Euromedex, 2000-C) for 10 min at 37°C, washed in PBS and permeabilized in 0.01% Triton X-100 (Euromedex, T8787) in PBS (PBT) at room temperature before being placed in blocking solution (PBT with 3% BSA) at 4°C for 4 h. Primary antibodies are applied in blocking solution at 4°C overnight. After washes in PBT at room temperature, embryos are incubated with secondary antibodies, DAPI and/or phalloidin at room temperature for 1 h. Embryos are washed in PBS and imaged immediately after.

| Primary antibodies | Dilution | Provider |
| --- | --- | --- |
| Atp1a | 1:100 | Santa Cruz Biotechnology, sc-21712 |
| Aqp3 | 1:500 | Novus Biologicals, NBP2-33872 |
| Cdh1 | 1:500 | eBioscience, 14-3249-82 |
| Ctnnb1 | 1:100 | Cell Signaling Technology, 8480 |
| Pard6B | 1:50 | Santa Cruz Biotechnology, sc-166405 |
| Prkcz | 1:50 | Santa Cruz Biotechnology, sc-17781 |

| Secondary antibody and dyes | Dilution | Provider |
| --- | --- | --- |
| Alexa Fluor Plus 488 anti-mouse | 1:200 | Invitrogen, A32723 |
| Alexa Fluor 546 anti-mouse | 1:200 | Invitrogen, A11003 |
| Alexa Fluor Plus 488 anti-rabbit | 1:200 | Invitrogen, A32731 |
| Alexa Fluor 488 anti-rat | 1:200 | Invitrogen, A11006 |
| Alexa Fluor 546 anti-rat | 1:200 | Invitrogen, A11081 |
| Alexa Fluor 633 anti-rat | 1:200 | Invitrogen, A21094 |
| Alexa Fluor 633 phalloidin | 1:200 | Invitrogen, A22284 |
| DAPI | 1:1000 | Invitrogen, D1306 |

#### Micropipette aspiration

As described previously (*26, 27*), a microforged micropipette coupled to a microfluidic pump (Fluigent, MFCS EZ) is used to measure the surface tension of embryos. In brief, micropipettes of radii 25.5-31.5 *µ*m are used to apply step-wise increasing pressures against the TE lining the blastocoel until reaching a deformation which has the radius of the micropipette (*Rp*). At steady-state, the surface tension *γ* of the blastocyst is calculated from the Young-Laplace’s law applied between the embryo and the micropipette: *γ* = *P*_*c*_/2 (1 */R_p_* - 1/*R*), where *P*_*c*_ is the critical pressure used to deform the embryo of radius *R.* Based on this measurement, we then use the Young-Laplace law between the blastocyst and the outside medium to calculate the hydrostatic pressure *P* of the embryo: *P* = 2 *γ / R.*

### Microscopy

Live imaging is performed using an inverted Zeiss Observer Z1 microscope with a CSU-X1 spinning disc unit (Yokogawa). Excitation is achieved using 488 and 561 nm laser lines through a 63x/1.2 C Apo Korr water immersion objective. Emission is collected through 525/50 and 595/50 band pass filters onto an ORCA-Flash 4.0 camera (C11440, Hamamatsu). The microscope is equipped with an incubation chamber to keep the sample at 37°C and supply the atmosphere with 5% CO_2_.

Surface tension measurements are performed on the same microscope using a 40x/0.95 Plan Apo Korr dry objective without CO2 supply in FHM medium.

Immunostainings are imaged on the same microscope using 405, 488, 561 and 642 nm laser lines through a 63x/1.2 C Apo Korr water immersion objective, 450/50 nm, 525/50 nm, 595/50 band pass or 610 nm low pass filters. Alternatively, an inverted Nikon Eclipse-Ti microscope with a CSU-X1 spinning disc unit (Yokogawa) is used. Excitation is achieved using 405, 488, 561 and 642 nm laser lines through a 60x/1.4 oil immersion DIC N2 PL APO VC objective. Emission is collected through 525/50 nm, 605/40 nm, 629/62 nm band pass or 640 nm low pass filters onto a CoolSnap HQ2 camera. Otherwise, a scanning confocal Zeiss LSM700 equipped with 405,488, 555 and 639 nm laser lines is used. Acquisition through a 63x/1,4 OIL DICII PL APO objective onto PMTs.

High temporal resolution imaging is performed using a Viventis Microscopy LS1 Live light-sheet microscope. Fluorescence excitation was achieved with a dual illumination scanned Gaussian beam light sheet of ∼1.1 *µ*m full width at half maximum using 488 and 561 nm lasers. Signal was collected with a Nikon CFl75 Apo LWD 25x/1.1 objective and through 525/50-25 or 561/25 band pass filter onto an Andor Zyla 4.2 Plus sCMOS camera. The microscope is equipped with an incubation chamber to keep the sample at 37°C and supply the atmosphere with 5% CO_2_.

### Data analysis

#### Microlumens area

Using FIJI (*43*), we track a given cell-cell contact over time and measure all of its microlumens by identifying their equatorial plane in 3D. We draw the outline of all individual microlumens and measure in 2D their cross-sectional area over time. Using R, we fit the dynamics of microlumens area with a piece-wise linear regression considering a single inversion point (*44, 45*). This delivers two slopes giving the swelling and discharge rates as well as the inversion times for a given contact.

Cells are determined to be TE or ICM based on the presence or absence of contact to the outside medium respectively. Interfaces are classified as a “near” or “away” from the final position of the blastocoel by locating the blastocoel and bisecting the embryo in two halves, one with and the other without the blastocoel in it.

#### Radius of curvature

Using FIJI, we select a cell interface that is in the equatorial plane of two contacting cells. We manually fit a circle onto the interface or onto either sides of the microlumen. For the curvature of cell-cell interfaces, we measure all interfaces that can be measured in fixed embryos with (+) or (-) Blebbistati.nFor microlumen measurements, we measure all microlumens that are large enough to be measured 60 min after the time of inversion. Images and ROIs are provided.

Cells are determined to be TE or ICM based on the presence or absence of contact to the outside medium respectively. Interfaces are classified as a “near” or “away” from the final position of the blastocoel by locating the blastocoel and bisecting the embryo in two halves, one with and the other without the blastocoel in it.

#### Blastocoel location in chimeric embryos

Cells from chimeric embryos are classified as ICM when they have no contact to the outside medium, as mural TE when they have a contact with the outside medium and with the blastocoel and as polar TE when they have a contact with the outside medium and no contact with the blastocoel.

The number of cells from either donor embryo is counted for each tissue (ICM, polar and mural TEs). We calculate the relative proportion of each donor embryo to each tissue and the ratio of mural TE to the total TE for a given donor embryo.

#### Apical/basal intensity ratio

Using FIJI, we select confocal slices cutting through the equatorial plane of a TE cell lining the blastocoel. We draw a ∼1 *µ*m thick segmented line along the cell-medium interface and measure the mean apical intensity. We draw a line spanning the cell-cell contacts and blastocoel interface and measure the mean basal intensity. We then calculate the apical to basal ratio. We measured 3 cells per embryo. Images and ROIs are provided.

#### Variability ratio

Using FIJI, we select confocal slices cutting through the equatorial plane of a cell. We measure 90 min before the appearance of the first microlumens and 90 min after. Using a 1 *µm* thick segmented line, we draw along the cell-cell contact and we measure the mean and standard deviation of the intensity along this contact. The coefficient of variation is calculated by dividing the standard deviation by the mean intensity. The variability ratio is determined by dividing the coefficient of variation before and after the appearance of microlumens.

For embryos that do not form a blastocoel (when treated with Y27632 or sucrose), the latest time of appearance of microlumens is determined in control embryos from the same experiment. Measurements are then performed 90 min before and 90 min after this time. Images and ROIs are provided.

Cells are determined to be TE or ICM based on the presence or absence of contact to the outside medium respectively. Interfaces are classified as a “near” or “away” from the final position of the blastocoel by locating the blastocoel and bisecting the embryo in two halves.

#### Statistics

Mean, standard deviation, minimum, maximum, lower and upper quartiles, median, paired and two-tailed Student’s *t-test,* single-tailed Mann-Whitney *U-test* and Chi-squared *p* values are calculated using Excel (Microsoft) or R (R Foundation for Statistical Computing). Statistical significance is considered when *p <* 10^−2^ Whisker plots show the minima, lower quartile, median, upper quartile and maxima.

The sample size was not predetermined and simply results from the repetition of experiments. No sample was excluded. No randomization method was used. The investigators were not blinded during experiments.

#### Code availability

The code of the simulations is available on the following repository under a MIT license: https://github.comN irtualEmbryo/lumen_network

## Supplementary Text

In this Supplementary Text we detail the equations of the physical model of the network of microlumens and the numerical implementation of the simulations. Lumens are considered in 2 dimensions, and are supposed of symmetric shape (see Supplementary Text Fig.[1]). They are connected by pipes of diameter *d,* representing the intercellular space between two adhesive membranes, which allows them to exchange fluid. The diameter *d* ∼ *20nm* is considered very small compared to lumen curvature radii and pipe length of *100nm* to *µm* sizes.

## 1 Area dynamics for two lumens

We consider first the case of two connected lumens. Each lumen i is characterized by its radius of curvature *R*_*i*_, its lineic tension γ_*i*_ and pressure *P*_*i*_. Force balance takes the form of Laplace’s law

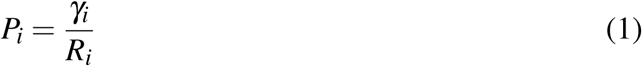

and by tension balance at the contact point

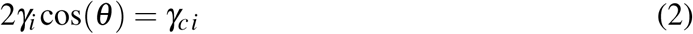

where γ_*c,i*_ is the surface tension at cell-cell contact for lumen *i.*

### 1.1 Laplace’s pressure in one lumen

The pressure inside each lumen *i* is calculated as function of its area by geometrical considerations, using the contact angle θ_*i*_ (see Supplementary Text Fig.[1]). The area *A*_*i*_ of the lumen *i* can be decomposed as

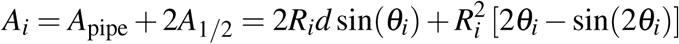

**Supplementary Text Figure 1:**
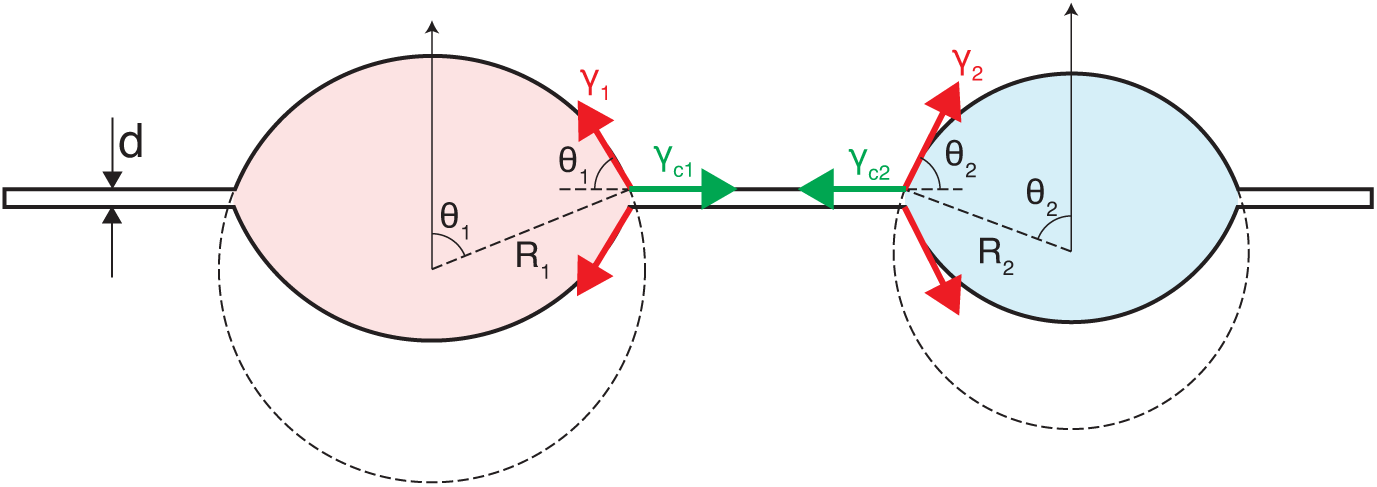
Geometrical parametrization of two lumens exhanching fluid via a pipe of diameter *d.* Each lumen is symmetric, with radius of curvature *R*_*i*_ (*i* = 1, 2) and contact angle *θ_i_.* The lineic tensions γ_*i*_ and contact tensions γ_*c,i*_ are shown so that Eq.(2) is satisfied.

Assuming *d* ≪ *R*_*i*_ we find

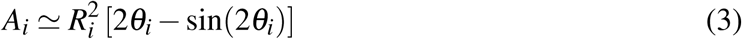

and it yields

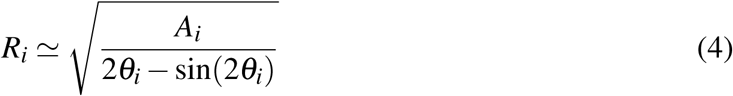

hence, using Eq. (1), we find the pressure of the lumen to be

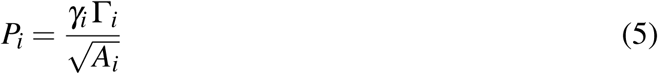

where 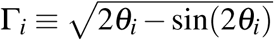.

### 1.2 Hydrodynamic flux and phase diagram

Two lumens with different curvatures and/or tensions will have different Laplace’s pressures, which gives rise to a pressure gradient and hence to a flow of fluid from the lumen with higher pressure to the one with a lower pressure. If we denote the two lumens 1 and 2, and assuming mass conservation *A*_tot_ = *A*_1_ + *A*_2_ = cte, the time evolution for the size of the lumens is given by the dynamical equation

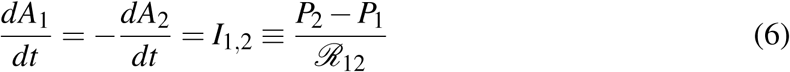

where we denoted *I*_1,2_ the flux from lumen 2 to 1 (*I*_2,1_ from 1 and 2). *ℛ*_12_ is the hydraulic resistance of the pipe, which is proportional to its length *𝓁*_12_

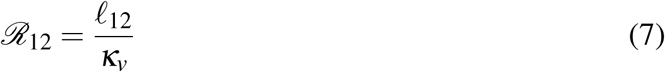

*κ_v_* is a friction coefficient given by Poiseuille’s law, that depends on the pipe width *d* and fluid viscosity *η*. For a 2-dimensional pipe, its expression is given by

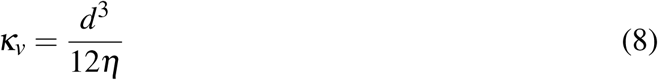

Eq. (6) is a deterministic equation, and for given initial conditions, we can predict which lumen between 1 and 2 will win, i.e. will grow, knowing the sign of the flux *I*_12_. The pressures *P*_*i*_ are given by Eq.(5) Thus, if *P*_2_ *> P*_1_, the lumen 1 grows. Expressing the flux *I*_12_ as function of the parameters *γ_i_ γ_ci_, A_i_; i* = 1, 2, we can build a phase diagram for the direction of the flux is summarized in Supplementary Text Fig.2.

### 1.3 Pumping

We now consider the case where the lumen is able to pump additional fluid through its interface. The flux *ℒ_i_* of pumped fluid is assumed to be proportional to the lumen perimeter

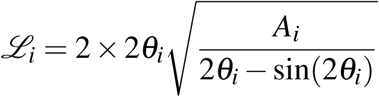

and to a pumping rate per unit length *λ* Here we do not consider pumping in the pipe itself. For an isolated lumen *i* with pumping, the time evolution of its size is given by

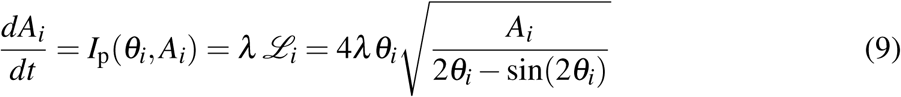

For the case of two connected lumens with the same pumping rate per unit length *λ*, their time evolutions are given by the equations

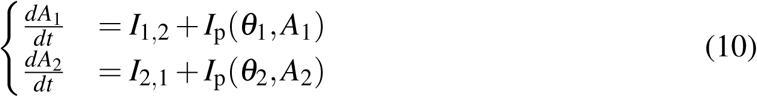

**Supplementary Text Figure 2:**
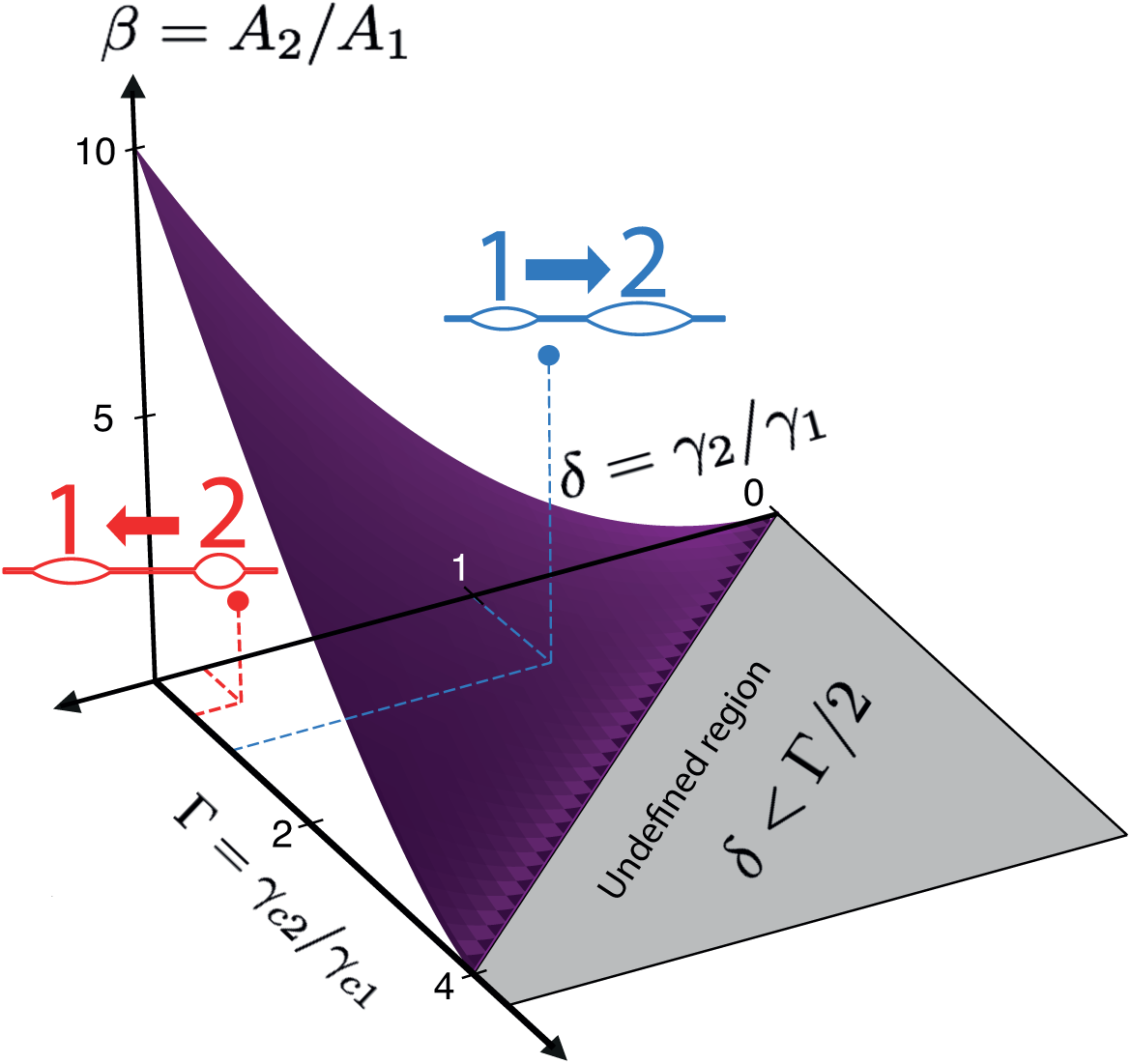
Phase diagram for the direction of the fluid flux between two lumens as function on the area asymmetry *β* = *A*_2_*/A*_1_, lumen tension asymmetry *δ* = *γ*_2_/*γ*_1_ and adhesive contact tension asymmetry Γ= *γ*_*c*2_/*γ*_*c*1._

## 2 Network of lumens

### 2.1 Dynamical equations

We can easily generalize Eq. (10) for a network of lumens *i* = l, 2, The network is described as an undirected graph *G* = {*V, E*}, where *V* is a set of vertices representing the lumens *i* = 1, 2, … and *E* is a set of edges (*i, j*) ∈ *V* x *V* representing the pipes. The dynamics is described by a set of coupled ordinary equations *i* = l, 2, …

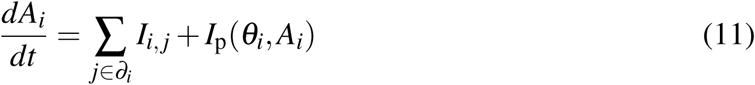

with *∂_j_* is the set of neighbors of the lumen *i.*

### 2.2 Numerical simulations

#### 2.2.1 Graph generation

We generate the lumen network as a set of connected vertices. Each vertex is a lumen with coordinates (*x,y*), connected to its neighbors. Self-loops are not allowed. The coordinates of the points are saved in a file (*lumen_coord.dat*) for graphic representation. The topology of the network can be chosen arbitrarily, but it is restricted here to hexagonal or triangular lattices. To each point is associated an initial area *A,* drawn in a truncated normal distribution *N*(*µ, σ*)_*A>P*_, such that *p* = 0. l. The parameters of the normal distribution are chosen as *µ* = l and *σ* = 0.1 here, and we verified that the results do not change qualitatively with this particular choice. To each lumen *i* is associated lineic lumen (*γ_i_*) and contact (*γ_c,i_*) tensions, that set the lumen contact angle *θ_i_.* The structure of the graph (set of edges) is saved in a file (*lumen _lumen.dat*), with the length of each edge *𝓁_ij_,* determining the hydraulic resistances. The initial parameters of the lumen (set of vertices) are saved in another file (*lumen.dat*) to allow averaging over the same structures. Other files are needed for the algorithm to process, such as a configuration file (*test.ini*) that sums up the input parameters, such as the tube diameter (we chose here *d* = 0.01). We call *regular* a lattice where vertices are equidistant from each others (usually, the length is set to 1), imposing hence the same resistance for every edge (*i,* j).

#### 2.2.2 Bridges

When the network is evolving, some lumens will shrink and their area will tend to zero. To avoid divergence, we consider a lumen *i* as empty when *A*_*i*_ < *d*^2^. An empty lumen cannot grow anymore but can still let the fluid pass through. If the empty lumen connects two lumens only, it is deleted from the network, and the distance between the neighbors is calculated summing the two initial pipe lengths (e.g. if lumen 2 connects 1 and 3 but disappears, 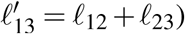. If the empty lumen has only one neighbor, it simply disappears from the graph. If the disappearing lumen is connected to more than two lumens, it will be replaced by a *bridge.* A bridge has an area *A*_*b*_ equal to zero and simply let fluid pass through. Its pressure *P*_*b*_ is calculated from mass conservation, stating that the sum of incoming fluxes should vanish exactly (Kirchhoff’s law):

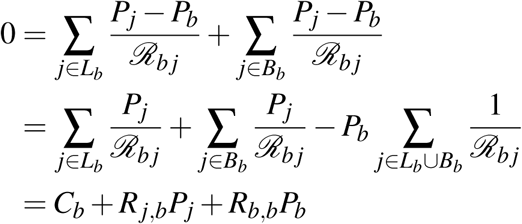

where *L*_*b*_ and *B*_*b*_ designate the set of neighboring lumens and bridges connected to a bridge *b.* Considering all bridge pressures, collected within a vector 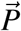, the coupled equations above yield a linear system

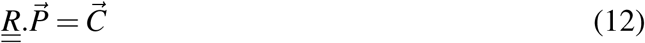

The vector 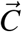 collects the pressures of the lumens weighted by the hydraulic resistances of pipes linking bridges with their neighboring lumens, and the matrix 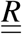. is a matrix with coefficients (*r*_*i,j*_) defined by *r*_*i,j*_ = **1**/*ℛ* _*i,j*_ and 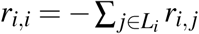.

The two types of connections to a bridge, i.e. *bridge-lumen* and *bridge-bridge,* are stored in specific data files (*bridge_lumen.dat* and *bridge_bridge.dat* respectively). As the network evolves, the number of lumens will decrease, reducing the computational cost of the simulation for large networks.

#### 2.2.3 Numerical scheme

The numerical integration of the set of equations Eq. (11) is done using the library **scipy.integrate** from Python2.7, more precisely the method *odeint*^1^. Since the method does not provide an event handle to detect if a lumen is empty, the numerical solving is done by restarting the integration for small time frames, checking the conditions and starting it again. However, because of the dynamics, the disappearance of lumens will change the network quickly at small times, and more slowly at larger times. Moreover, the dynamics at large times is much slower than at small times, because of the competition between larger lumens is slower. We therefore implemented an *adaptive time stepping,* that depends on the number of lumens *N* as

**Supplementary Text Figure 3:**
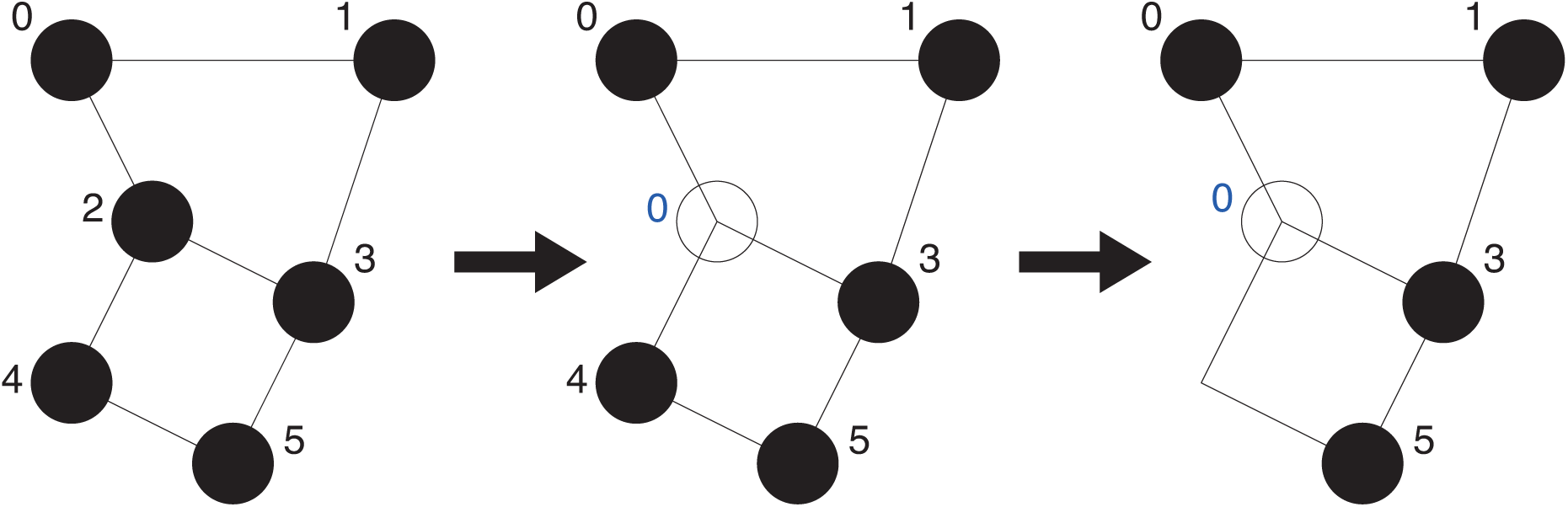
A simple network evolution. Lumen 2 becomes bridge 0, then lumen 4 disappears but the link between bridge 0 and lumen 5 still exists.

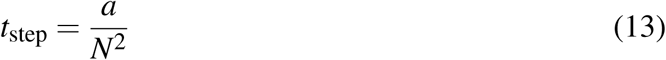

where *a* = 0.1 is an arbitrary constant, chosen to obtain a smooth dynamics. The time frame for which the integration proceeds before conditions are checked again is set as *t*_frame_= *k*.*t*_step,_ with *k* = 20 an arbitrary constant.

The calculation of the area change 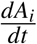 is split into four steps:

1. the flux through between lumens *i* and 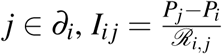 is calculated, adding 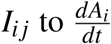, subtracting 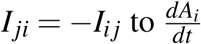.
2. the bridge pressures are calculated by solving the linear system (12) using the library **numpy.linalg,** with the method **solve.**
3. similar to 1., the fluxes through bridge-lumen connections are calculated, 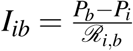.
4. if pumping is included and the lumen is not empty, then the area change for lumen *i* is calculated using Eq. (9).

In the end, the area change for lumen *i* is given by

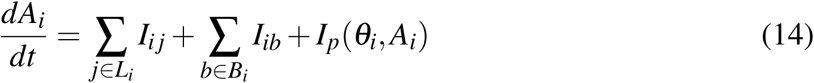

**Supplementary Text Figure 4:**
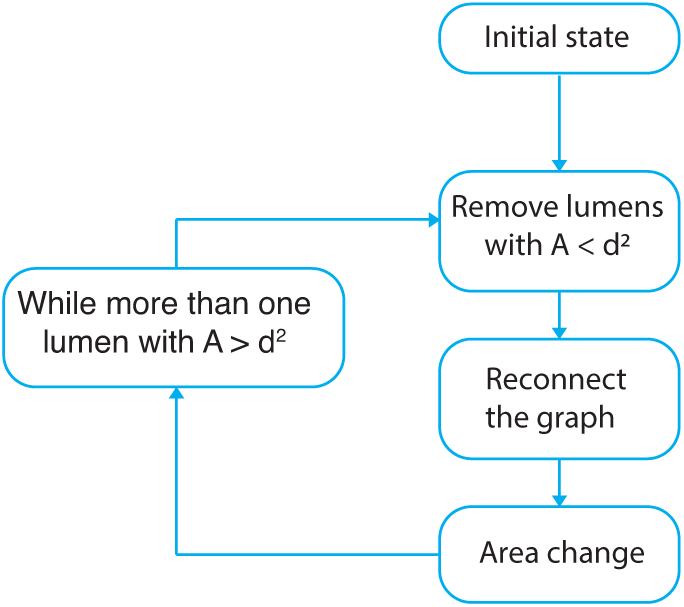
Basic steps of the script.

The termination condition is defined as **”There is only one non-empty lumen”.** Areas are stored in an output file (*area.dat*), and each time step is recorded. The area conservation is checked at every step, and errors are stored in a log file (*error.log*). The initial and final configuration are stored in a third file (*area.log*) in order to give an easy access to the important observables. A fourth file tracks the disappearance and relabeling of lumens into bridges if needed, as well as the reconnections in the graph.

## 3 Robustness of the simulations

In this section, we present additional results to show the robustness of our simulations. We will denote 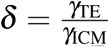 the tension asymmetry where γ_*T E*_ (γ_*ICM*_) is the tension of a lumen at the TE-ICM interface (ICM-ICM interface).

### 3.1 Effect of the initial area distribution

The initial area distribution of the lumens is a truncated Gaussian distribution

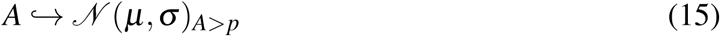

such that *µ* = 1. *p* = 0.1 is an arbitrary constant and minimal initial area allowed. The width of the distribution is usually taken to be *σ* = 0.1. Supplementary Text Fig. 5 shows the effect of broader distributions on the probability for the final lumen to be at the TE-ICM interface: as the width of the area distribution increases, the sharpness of the transition decreases. This can be simply interpreted as the consequence of an increased probability of starting with a large initial lumen in a region of larger tension, the size asymmetry compensating the imposed tension asymmetry.

**Supplementary Text Figure 5:**
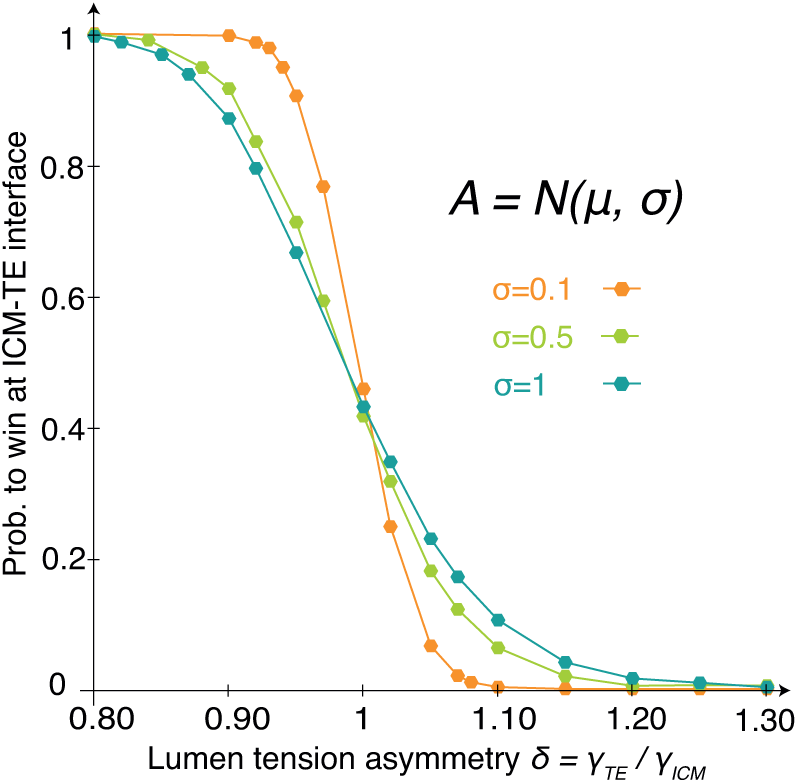
Winning probability for lumens at the TE-ICM interface (external layer) for a regular hexagonal lattice with 2 layers, for different widths of the initial areas distribution. The initial areas are drawn from a truncated normal law (A > 0.1) with the same average (µ = 1) but different standard deviations (σ= 0.1, 0.5, 1). The lumen network is the regular network similar to the one used in Fig. 3C. Each point results from at least 5000 simulation s

### 3.2 Noisy lattices

Networks are generated from regular lattices, with identical distances between lumens. *Noisy* lattices are generated from a regular lattice, by displacing randomly the positions of the vertices by a vector 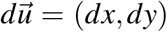 of components 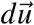 drawn from a normal distribution 𝒩 (0, 0.05). For a given lumen network, multiple configurations are averaged with the same number of simulations each. Noisy lattices show no deviations from the winning probability distribution of regular lattice, for both the tension asymmetry ratio δ and contact tension asymmetry ratio г (see Supplementary Text Fig. 6)

**Supplementary Text Figure 6:**
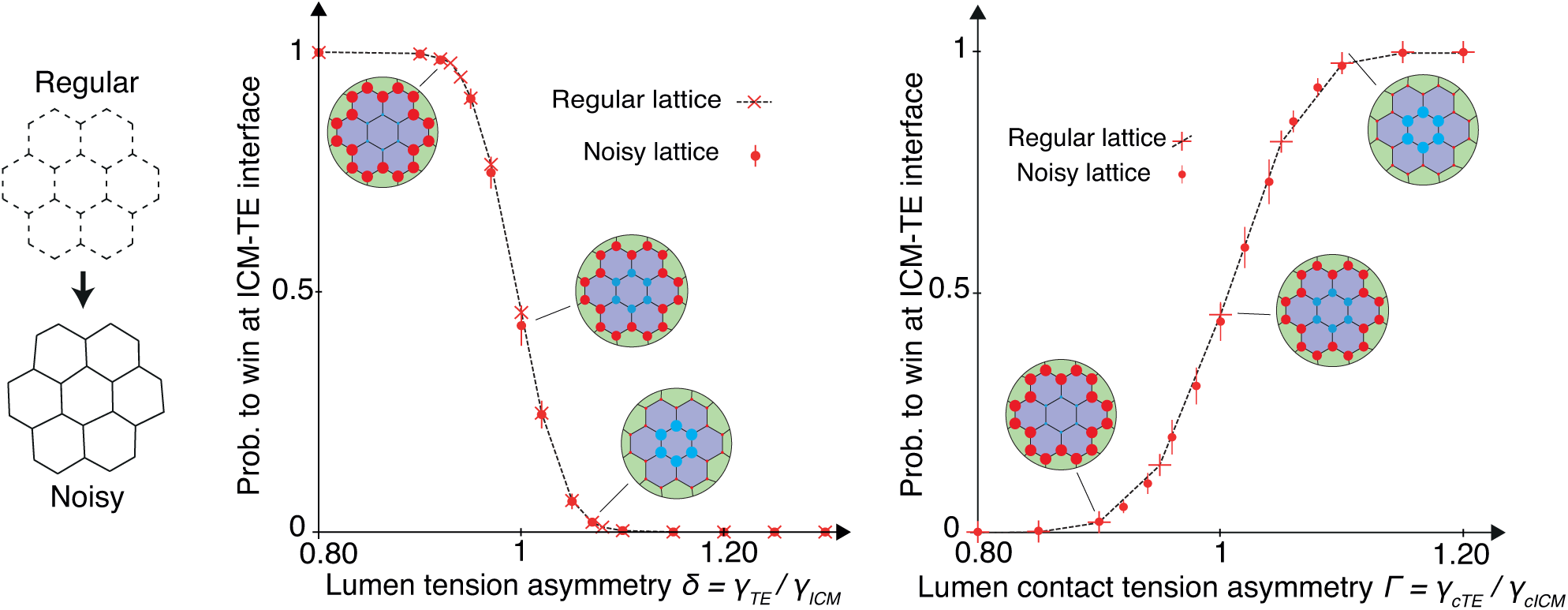
(Left) A hexagonal noisy lattice (bottom), calculated from the regular hexaongal (top) lattice. (Center and right) Winning probability for lumens at the TE-ICM interface (external layer) for regular(red crosses, dashed lines) and noisy (red dots, red vertical bar for standard deviation) hexagonal lattices, as functions of the tension asymmetry *δ* (center) and the contact tension asymmetry (right). Each point of the regular lattice results from at least 2000 simulations, the noisy lattice is the average of 25 configurations of the noisy lattice, 2000 simulations each.

### 3.3 Variations in topology

We also tested two different topologies: hexagonal and triangular lattices. Hexagonal lattices are preferred as they more faithfully represent the connection between cells in 2 dimensions, corresponding essentially to tricellular junctions. Triangular lattices are on the contrary more akin to an isotropic network. The TE-ICM interface is defined as the outer layer, with *γ_JcM_* = 1 and *γ_TE_* changed to tune the tension asymmetry *δ.* The topology or size of the network do not affect the qualitative behavior of the curve. However, an isotropic topology favors generally lumens inside the network and increasing the size of the network also marginally displaces the curve towards lower *δ* (see Supplementary Text Fig.7). Hence as the network grows in size, we expect a higher asymmetry to be necessary to statistically direct the final lumen towards the network boundary.

**Supplementary Text Figure 6:**
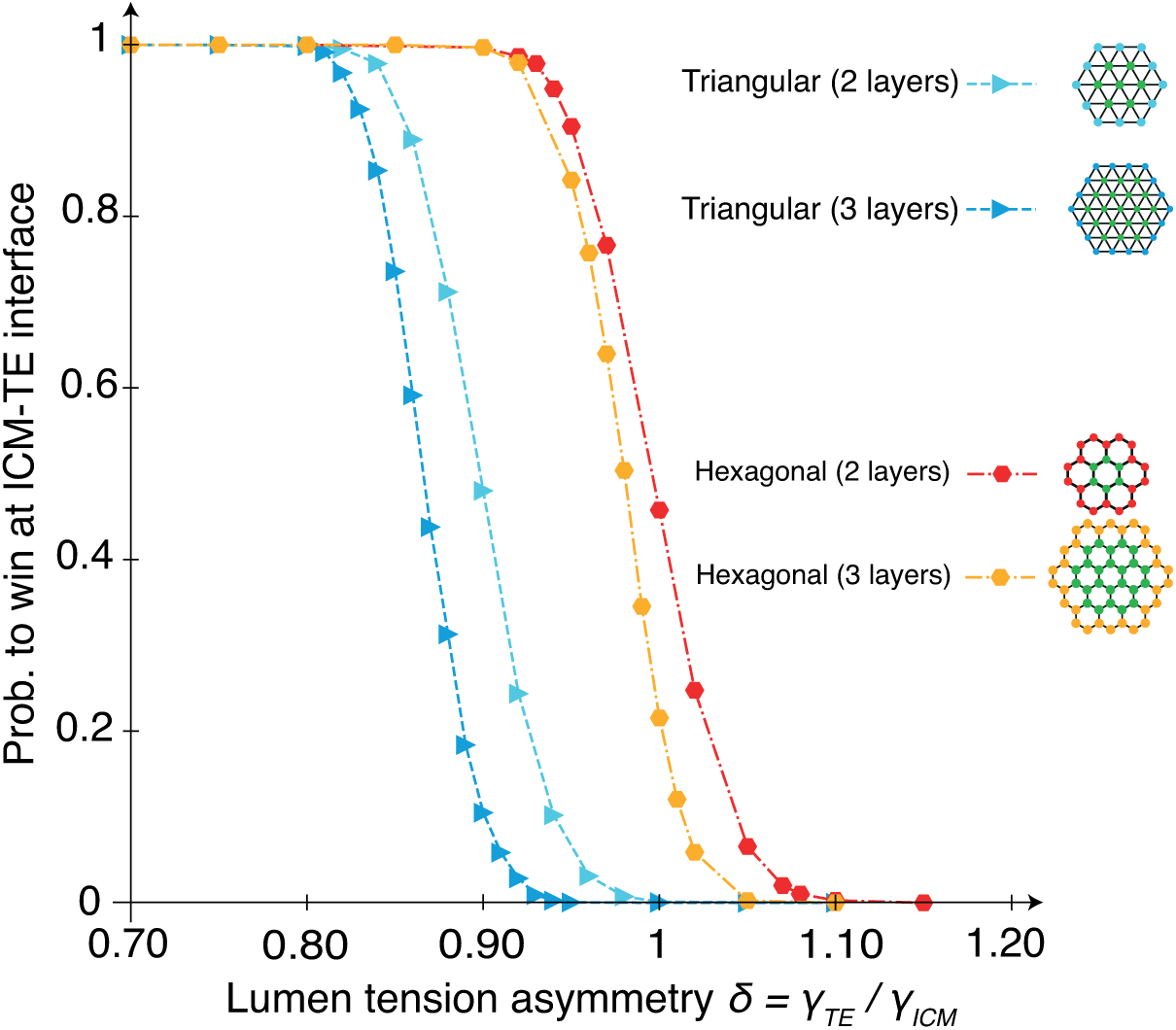
Winning probability for lumens at the TE-ICM interface (external layer) for regular hexagonal and regular triangular lattices of different sizes. An inset shows the lattices with lumens (green for ICM-ICM, colored for ICM-TE). Each point results from at least 2000 simulations.

## Footnotes

1 See documentation

